# Spatial charting of single cell transcriptomes in tissues

**DOI:** 10.1101/2021.11.24.469915

**Authors:** Runmin Wei, Siyuan He, Shanshan Bai, Emi Sei, Min Hu, Alastair Thompson, Ken Chen, Savitri Krishnamurthy, Nicholas E. Navin

## Abstract

Single cell RNA sequencing (scRNA-seq) methods can profile the transcriptomes of single cells but cannot preserve spatial information. Conversely, spatial transcriptomics (ST) assays can profile spatial regions in tissue sections, but do not have single cell genomic resolution. Here, we developed a computational approach called CellTrek that combines these two datasets to achieve single cell spatial mapping. We benchmarked CellTrek using a simulation study and two *in situ* datasets. We then applied CellTrek to reconstruct cellular spatial structures in existing datasets from normal mouse brain and kidney tissues. We also performed scRNA-seq and ST experiments on two ductal carcinoma *in situ* (DCIS) tissues and applied CellTrek to identify tumor subclones that were restricted to different ducts, and specific T cell states adjacent to the tumor areas. Our data shows that CellTrek can accurately map single cells in diverse tissue types to resolve their spatial organization.

## Introduction

Single cell RNA sequencing (scRNA-seq) methods have greatly expanded our understanding of the gene expression programs of diverse cell types and their role in development and disease^1-5^. However, scRNA-seq inherently loose cellular spatial information during the tissue dissociation step, which is critical for understanding cellular microenvironment and cell-cell interactions^6-8^. While spatial sequencing methods, including spatial transcriptomics (ST)^9^ and Slide-seq^10^, can spatially profile gene expression across tissue sections, they are limited to measuring small regions with mixtures of cells and cannot easily provide single cell information. To address this issue, computational approaches (e.g., cell2location, RCTD) have been designed to deconvolute ST spots into proportions of different cell types^11-17^. However, spatial deconvolution methods are limited to inferring only cell type proportions for each spot, and cannot achieve single cell resolution. Additionally, deconvolution methods have limited ability to further resolve cell types into more granular “cell states” (expression programs) that reflect different biological functions. Finally, most deconvolution methods can only predict categorical labels and cannot infer continuous cell information (e.g., lineage trajectories, gene signatures, continuous phenotypes) at a spatial resolution.

Here we introduce CellTrek, a computational toolkit that can directly map single cells back to their spatial coordinates in tissue sections based on scRNA-seq and ST data. This method provides a new paradigm that is distinct from ST deconvolution, enabling a more flexible and direct investigation of single cell data with spatial topography. The CellTrek toolkit also provides two downstream analysis modules, including SColoc for spatial colocalization analysis and SCoexp for spatial co-expression analysis. We benchmarked CellTrek using simulations and *in situ* datasets. We then applied CellTrek to existing datasets from normal mouse brain^18^ and kidney^19^ tissues as well as data that we generated from two human ductal carcinoma *in situ* (DCIS) samples to study the organization of cell types/states at single cell spatial resolution.

## Results

### Overview of CellTrek toolkit

CellTrek first integrates and co-embeds ST and scRNA-seq data into a shared feature space (Fig. 1, Methods). Using the ST data, CellTrek trains a multivariate random forests (RF) model^20^ to predict the spatial coordinates using shared dimension reduction features. A spatial non-linear interpolation on ST data is introduced to augment the spatial resolution. The trained model is then applied to the co-embedded data to derive an RF-distance matrix which measures the expression similarities between ST spots and single cells supervised by spatial coordinates. Based on the RF-distance matrix, CellTrek produces a sparse spot-cell graph using mutual nearest neighbors (MNN) after thresholding. Finally, CellTrek transfers spatial coordinates for cells from their neighbor spots. To improve the compatibility, CellTrek can accept any cell-location probability/distance matrix calculated from other methods (e.g., novoSpaRc^21^) as an input for cell spatial charting. Additionally, we provide a graphical user interface (GUI) for interactive visualization of the resulted CellTrek map.

**Fig. 1.**
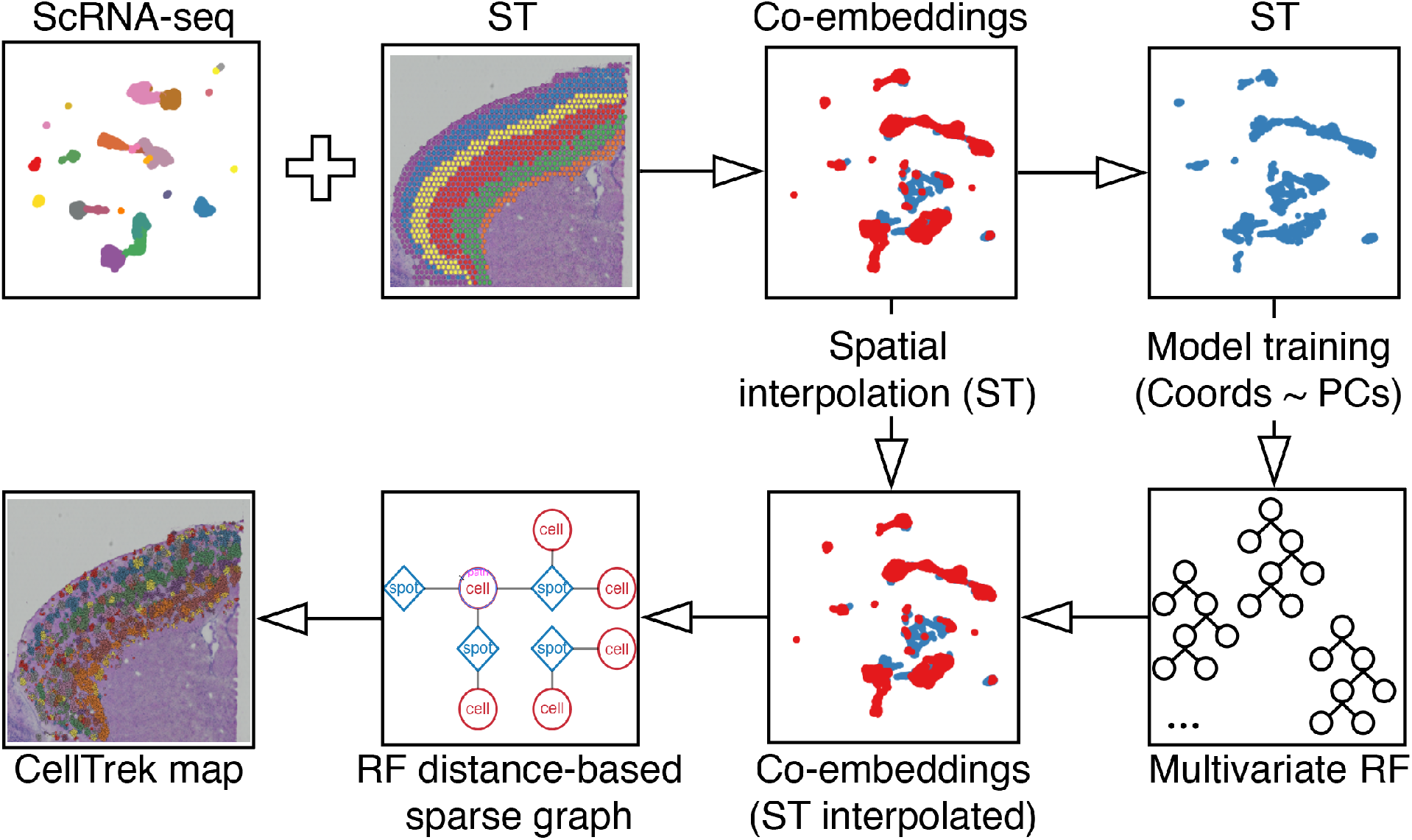
Overview of the CellTrek workflow. CellTrek first co-embeds scRNA-seq and ST datasets into a shared latent space. Using the ST data, CellTrek trains a multivariate random forests (RF) model with spatial coordinates as the outcome and latent features as the predictors. A 2D spatial interpolation on the ST data is introduced to augment the ST spots. The trained RF model is then applied to the co-embedded data (ST interpolated) to derive an RF-distance matrix which will be converted into a sparse graph using mutual nearest neighbors (MNN). Based on the sparse graph, CellTrek transfers the coordinates to single cells from their neighboring ST spots.

To recapitulate spatial relationships between different cell types, we developed a downstream computational module, SColoc, which summarizes the CellTrek result into a graph abstraction (Supplementary Fig. 1a, Methods). Three approaches, Kullback-Leibler divergence (KL), Delaunay triangulation (DT), and K-nearest neighbor distance (KD), are provided to calculate spatial dissimilarity between cell types. Based on the dissimilarity matrix, SColoc can construct a minimum spanning tree (MST) that represents a simplified spatial cellular proximity. The above steps will be iteratively executed on bootstrapping samples to generate consensus matrices (on dissimilarities or MSTs). Thereafter, a graph will be rendered through a GUI with tunable edge thresholding and color mapping functions. Additionally, SColoc provides a K-distance metric for measuring the spatial distance of cells to a selected reference group.

To investigate whether different expression programs are distributed across different topographic areas, we developed SCoexp which leverages the CellTrek coordinates to detect co-expression gene modules within the cells of interest (Supplementary Fig. 1b, Methods). First, SCoexp calculates a spatial kernel weight matrix based on their spatial distances. Using this weight matrix, SCoexp calculates spatial-weighted gene co-expression matrix. Thereafter, SCoexp utilizes consensus clustering^22^ (CC) or weighted correlation network analysis^23^ (WGCNA) to identify gene modules. For the identified modules, we can calculate module scores and investigate their spatial organizations.

### Benchmarking and simulations

To benchmark the performance of CellTrek, we exploited three spatial datasets, 1) a simulated scRNA-seq dataset with customized spatial patterns (Supplementary Fig. 2a, b); 2) a fluorescence *in situ* hybridization (FISH)-based single cell dataset of the *Drosophila* embryo^21^ (Supplementary Fig. 2d, e) and 3) a seqFISH dataset of mouse embryo^24^ (Supplementary Fig. 2g, h). We generated three corresponding ST datasets with each spot aggregating the five spatially nearest cells (Supplementary Fig. 2c, f, i).

We applied CellTrek to the scRNA-seq and ST data to reconstruct their spatial cellular maps. We then compared CellTrek to two additional cell charting methods: 1) NVSP-CellTrek which uses a reference-based novoSpaRc^21^, a spatial reconstruction method, to calculate a cell-spatial probability matrix, then leverages CellTrek to produce a spatial map, and 2) Seurat^25^ coordinate transfer (SrtCT) which uses the data transfer approach to transfer ST coordinates to single cells. Both CellTrek and NVSP-CellTrek reconstructed the original spatial pattern of the simulated data, while SrtCT only reconstructed a rough spatial relationship between cells and could not accurately map the cells (Supplementary Fig. 3a). Compared to NVSP-CellTrek, CellTrek mapped more cells with higher spatial density. To quantitatively evaluate these methods, we compared the spatial density of the cell charting results to the original spatial distribution across different cell types using the KL-divergence. Both CellTrek and NVSP-CellTrek achieved good performance with low KL-divergences, while SrtCT showed much higher discrepancies to the reference distribution (Supplementary Fig. 3b). In the *Drosophila* embryo data, CellTrek accurately reconstructed the original spatial layout with the lowest KL-divergences among three approaches (Supplementary Fig. 3c, d). We further investigated several known *Drosophila* embryogenic genes in the CellTrek results and found consistent spatial patterns to the previous study^21^(Supplementary Fig. 3e). In the mouse embryo data, we found that CellTrek and NVSP-CellTrek accurately reconstructed the original spatial structure, while CellTrek showed slightly higher KL-divergences in groups 5, 9 and 17 (Supplementary Fig.3 f, g). To investigate if CellTrek could reveal the developing spatial patterns of the mouse embryo, we selected a group of gut tube cells and found that there were spatial consistencies in some marker genes to the previous study^24^ (Supplementary Fig. 3h). We then performed a trajectory analysis using Monocle2^26, 27^ which showed that the pseudotime reflected the spatial developing pattern of the gut tube cells along with the anterior-posterior axis^24^ (Supplementary Fig. 3i).

We next assessed the performance of CellTrek on simulated data under three different simulation settings: 1) read counts, 2) spatial randomness, and 3) tissue densities. We evaluated CellTrek performance using KL-divergence and Pearson’s correlation on the cell spatial coordinates between the CellTrek map and the reference. Across the three simulations (with eight conditions each), CellTrek achieved good spatial reconstruction performances (Supplementary Fig. 4a,c,e,g,i) and showed lower KL-divergences and higher correlations compared to the permutation test (Supplementary Fig. 4b,f,j). However, increasing the spatial randomness will affect the performance of CellTrek and decrease the statistical significance (Supplementary Fig. 4f) while decreasing the read counts or the spot/cell density will result in sparse cellular maps (Supplementary Fig. 4a-b, g-j). Overall, this data suggests that CellTrek is a robust method for single cell spatial mapping under different experimental conditions.

### Topological organizations of mouse brain cells

We applied CellTrek to public mouse brain scRNA-seq (Smart-seq2)^18^ and ST datasets (Visium, 10X Genomics). We compared CellTrek to NVSP-CellTrek and SrtCT approaches. CellTrek reconstructed a clear layer structure of laminar excitatory neuron subtypes, in the order of L2/3 intratelencephalic (IT), L4, L5 IT, L6 IT, L6 corticothalamic (CT), and L6b, which matched to the cerebral cortex structure (Fig. 2a, Supplementary Data 1). NVSP-CellTrek showed a similar spatial layer trend thus demonstrating the flexibility and consistency of the CellTrek approach (Fig. 2a). However, NVSP-CellTrek resulted in a sparse cell mapping in some areas. SrtCT failed to accurately project cell locations to the histological image (Fig. 2a). We then employed Seurat label transfer (SrtLT) to predict the spatial distribution of each cell type as our reference^28^. KL-divergences between cell charting results and the reference suggested that CellTrek successfully recovered spatial cellular structures with the lowest KL-divergences among three approaches (Fig. 2b).

**Fig. 2.**
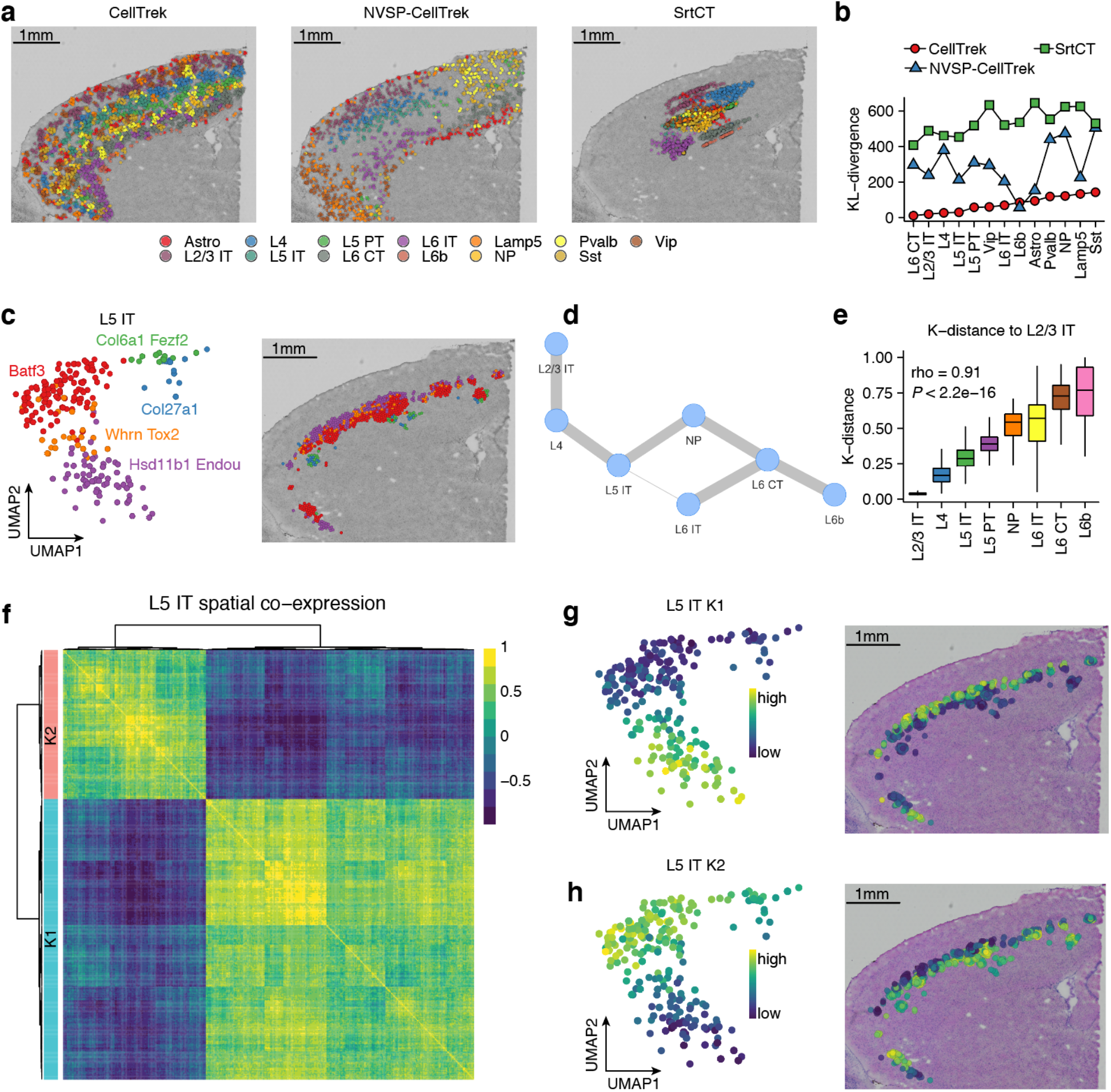
CellTrek reconstructs spatial organization in a mouse brain tissue. **a**, Comparison of CellTrek, NVSP-CellTrek and SrtCT results for single cell spatial charting in a mouse brain tissue. **b**, KL-divergence of spatial cell charting methods for each cell type using SrtLT as a reference. **c**, UMAP (left) and CellTrek map (right) of scRNA-seq data of L5 IT cell states. **d**, Spatial colocalization graph of glutamatergic neurons using SColoc. **e**, CellTrek-based spatial K-distance of glutamatergic neurons to L2/3 IT cells. Boxplots show the median with interquartile ranges (25–75%); whiskers extend to 1.5X the interquartile range from the box. **f**, Spatial co-expression modules (K1 and K2) identified in L5 IT cells using SCoexp. **g-h**, UMAPs of L5 IT cells showing the K1 module activity scores (**g**) and the K2 module activity scores (**h**) and their corresponding CellTrek maps.

Next, we asked if CellTrek could further uncover topological patterns of cell states within the same cell type. For example, L5 IT cells contain five expression states and showed a continuous trend on the UMAP in the order of Hsd11b1-Endou, Whrn-Tox2, Batf3, Col6a1-Fezf2 and Col27a1 (Fig. 2c, left). The L5 IT CellTrek map discovered a refined sub-layer architecture (Fig. 2c, right) which is consistent with a previous study^29^. To summarize the cell spatial colocalizations, we applied the SColoc to the CellTrek result using KL-based MST consensus graph. Glutamatergic neuron cell types constructed a linear backbone of the graph in the order of the layer structures (Fig. 2d). Spatial K-distance to the L2/3 IT cells showed a significant increasing trend in the same order of the graph (Spearman’s rho = 0.91, *P* < 2.2e-16) (Fig. 2e).

We then investigated how genes were spatially co-expressed within L5 IT cells using SCoexp. Two co-expression modules (K1, K2) were identified and showed different enrichments of biological functions (Fig. 2f, Supplementary Fig. 5a, b). The K1 module was highly active in cell states Hsd11b1-Endou, Whrn-Tox2 and spatially located in the outer layer, while the K2 module was highly active in Col27a1, Col6a1-Fezf2, and Batf3 (Fig. 2 c, g, h) and mainly located in the inner layer (Fig. 2 g, h). These results show that SCoexp is able to identify subtle transcriptional differences within the same cell type and infer their topological heterogeneity.

### Spatial cell charting of the mouse hippocampus

We also applied CellTrek to Slide-seq v2^30^ and scRNA-seq data^31^ from the mouse hippocampus. Unsupervised clustering of the Slide-seq data identified 12 clusters (G01-G12) with a highly organized spatial structure (Supplementary Fig. 6a). CellTrek mapped single cells to their spatial locations (Supplementary Fig. 6b), which is consistent with the Slide-seq clusters. Notably, the G06 matched with the Cornu Ammonis (CA) areas (Supplementary Fig. 6c), while CellTrek revealed a sequential mapping of the CA1, CA2, and CA3 principal cells that were not resolved by the Slide-seq clustering alone (Supplementary Fig. 6d). These results show that CellTrek can be applied broadly to different spatial genomic platforms, to achieve a more refined spatial cellular resolution.

### Spatial reconstruction of a mouse kidney tissue

We applied CellTrek to a public mouse kidney data^32^ and compared it to NVSP-CellTrek and SrtCT. CellTrek accurately reconstructed cellular spatial structures with distinct cell types located in different histological zones (e.g., cortex, outer medulla and inner medulla) (Fig. 3a, Supplementary Data 2). NVSP-CellTrek showed similar spatial patterns compared to CellTrek while SrtCT could not reconstruct accurate spatial organizations of the mouse kidney cells (Fig. 3a). Using SrtLT as a reference, both CellTrek and NVSP-CellTrek achieved overall low KL-divergence and NVSP-CellTrek showed higher KL-divergence for VSMC and RenaCorp cells (Fig. 3b). SrtCT showed the highest KL-divergence to the reference distribution. To further study the spatial cell expression dynamics, we inferred the trajectories of ProxTub and DistTub cells respectively and spatially mapped their pseudotime based on CellTrek. For ProxTub cells, we observed a continuous spatial trajectory that started from the outer part of the cortex to the inner part (Fig. 3c). This continuous anatomic change of ProxTub cells is consistent with previous studies^33, 34^. Similarly, DistTub cells also showed a continuous trajectory with a clear spatial pattern (Fig. 3d). Collectively, these results show that CellTrek can resolve the topological arrangements of continuous expression programs of single cells in tissues.

**Fig. 3.**
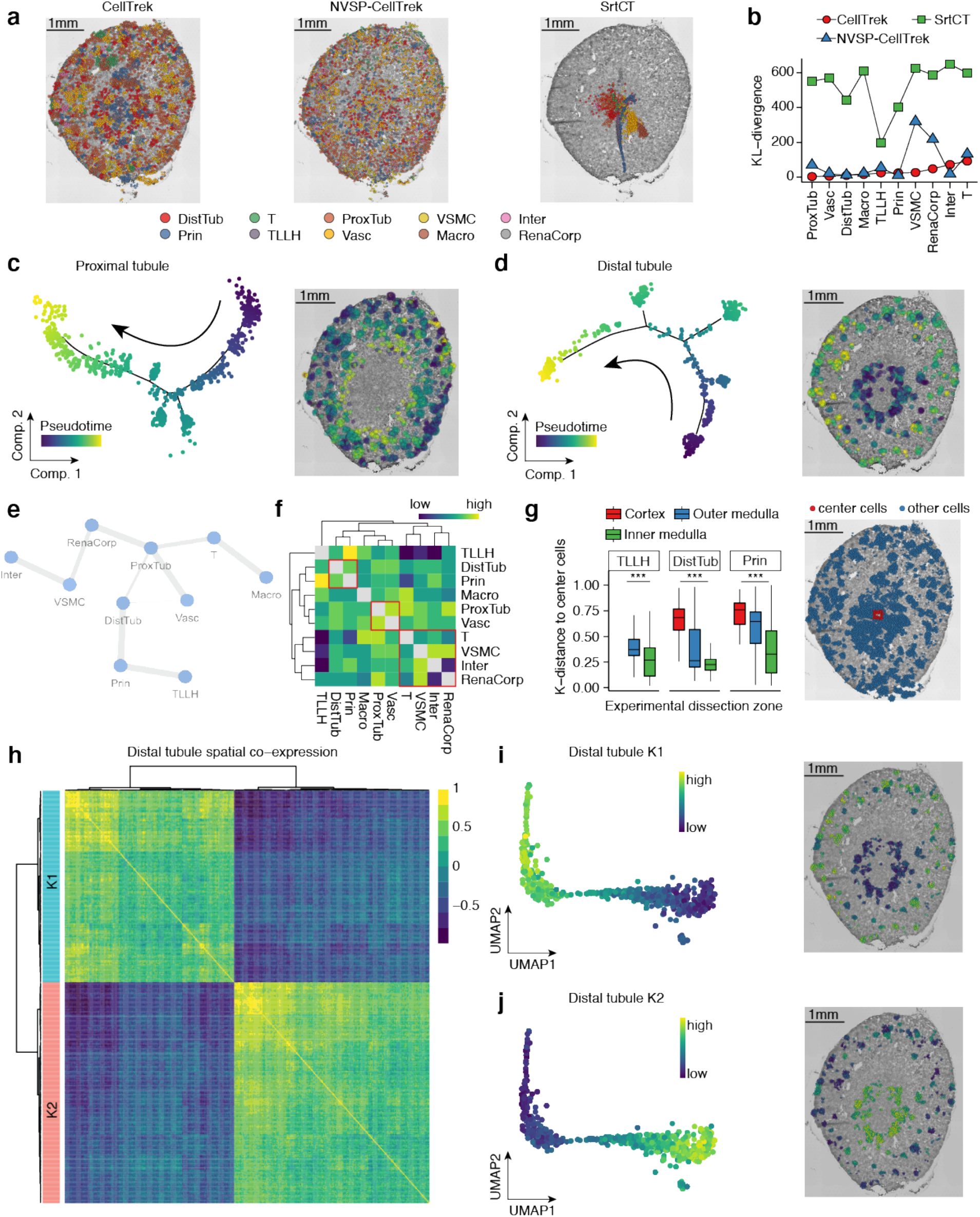
CellTrek reconstructs spatial organization in a mouse kidney tissue. **a**, Comparison of CellTrek, NVSP-CellTrek and SrtCT results for single cell spatial charting in a mouse kidney tissue. (DistTub: distal tubule cells, T: T cells, ProxTub: proximal tubule cells, VSMC: vascular smooth muscle cells, Inter: intercalated cells, Prin: principal cells, TLLH: the loop of Henle, Vasc: vascular cells, Macro: macrophages, RenaCorp: renal corpuscle cells) **b**, KL-divergence of spatial cell charting methods for each cell type using SrtLT as a reference. **c**, Trajectory analysis for proximal tubule cells (left) and spatial mapping of the pseudotime values in the tissue section (right). **d**, Trajectory analysis for distal tubule cells (left) and spatial mapping of the pseudotime values in the tissue section (right). **e**, Spatial colocalization graph of different renal cell types using SColoc. **f**, Spatial consensus matrix of different renal cell types. **g**, CellTrek-based spatial K-distance of TLLH, DistTub and Prin cells to the tissue center cells across experimental zonal dissections (left). Center cells as reference are shown on the right panel. *** indicates *P* < 0.001. Boxplots show the median with interquartile ranges (25–75%); whiskers extend to 1.5X the interquartile range from the box. **h**, Spatial co-expression modules (K1 and K2) identified in distal tubule cells using SCoexp. **i-j**, UMAPs of distal tubule cells showing the K1 module activity scores (**i**) and the K2 module activity scores (**j**) and their corresponding CellTrek maps.

We next summarized a cell spatial graph using SColoc. ProxTub cells were identified as the hub and connected to the RenaCorp, DistTub and other cell types (Fig. 3e). The consensus heatmap and hierarchical clustering showed similar patterns to the graph abstraction (Fig. 3f). Since the scRNA-seq data were collected from different zonal microdissections of the mouse kidneys^32^, we asked if CellTrek could recapitulate the experimental zonal information without the prior knowledge. Based on the CellTrek result, we calculated the K-distance for TLLH, DistTub and Prin cells to a group of cells from the center region. A consistent trend was observed that K-distances decreased from cortex to outer medulla then to inner medulla, suggesting that CellTrek successfully revealed the zonal structure of the mouse kidney (Fig. 3g). Further, in the DistTub cells, we identified two distinct spatial co-expression modules (K1 and K2) using SCoexp (Fig. 3h). The K1 module was enriched with metabolic pathways, renal system development and highly correlated with some distal convoluted tubule (DCT) genes (e.g., *Wnk1* and *Slc12a3*)^35, 36^ (Supplementary Fig. 5c). In contrast, K2 was enriched with cell-matrix pathways, purine metabolic pathways and correlated with distal straight tubule (DST) canonical genes (e.g., *Slc12a1* and *Umod*)^36^ (Supplementary Fig. 5d). These two modules displayed distinct patterns on the UMAP and the CellTrek map. K1 was highly active in the cortex area, whereas K2 was active in the medulla, which are consistent with the anatomic localizations of DCT and DST (Fig. 3i, j).

We further asked if CellTrek could improve our understanding of cell-cell communications by leveraging the spatial information. We conducted a cell-cell interaction analysis on the scRNA-seq data using CellChat^37^ and used the SColoc graph (Fig. 3e) to filter cell-cell pairs by assuming that colocalized cells will have a higher chance to interact with each other. Compared to the raw CellChat results, which predicted many non-specific interactions with all cell types interacting with each other (Supplementary Fig. 5e), the spatial filtering provided a reduced set of interactions that were more concise and specific (Supplementary Fig. 5f). Importantly, we identified several interactions which have been reported previously, including *Vegfa* which is expressed by the ProxTub interacted with its receptors *Flt1* and *Kdr* that are expressed by Vasc (Supplementary Fig. 5g)^38-41^.

### Spatial subclone heterogeneity in a DCIS breast cancer

We applied 3’ scRNA-seq (10X Genomics) and ST (Visium, 10X Genomics) to a DCIS sample (DCIS1) to profile 6,828 single cells and 1,567 ST spots. For the scRNA-seq data, clustering and differential expression (DE) analyses identified 5 major cell types, including epithelial, endothelial, fibroblast, myeloid and natural killer (NK)/T cells (Supplementary Fig. 7a). We applied CopyKAT^42^ to infer copy number profiles from the scRNA-seq data. We observed some clonal copy number alterations (CNAs) across all tumor cells, including gains on chromosomes 3q (*PIK3CA*), 8q (*MYC*), and 19p (*STK11*) and losses on chromosomes 8p (*PPP2R2A*), 10q (*PTEN*) and 14q (*AKT1*) (Fig. 4a). UMAP and dbscan clustering of the CNA profiles identified three major tumor subclones (clone1-3) with some distinct alterations, including 17q (*ERBB2*) gain and 11q (*ATM*) loss in clone2 and clone3, 1q (*MDM4* and *EPHX1*) gain in clone2 and 6q (*FOXO3*) loss in clone3 (Fig. 4 a, b). Based on the consensus CNA profiles, we constructed a phylogenetic tree which showed that clone1 was an earlier subclone that diverged from the main lineage, followed by clone2 and clone3 (Fig. 4c). Notably, these three subclones displayed transcriptional heterogeneity (Supplementary Fig. 7b). Hallmark gene set enrichment analysis^43^ identified several common pathways across all three subclones including MYC targets, oxidative phosphorylation and DNA repair (Fig. 4d). We also identified subclonal-specific signatures, including estrogen response pathways enriched in clone2 and clone3, and interferon alpha/gamma response, coagulation and complement pathways enriched in clone2.

**Fig. 4.**
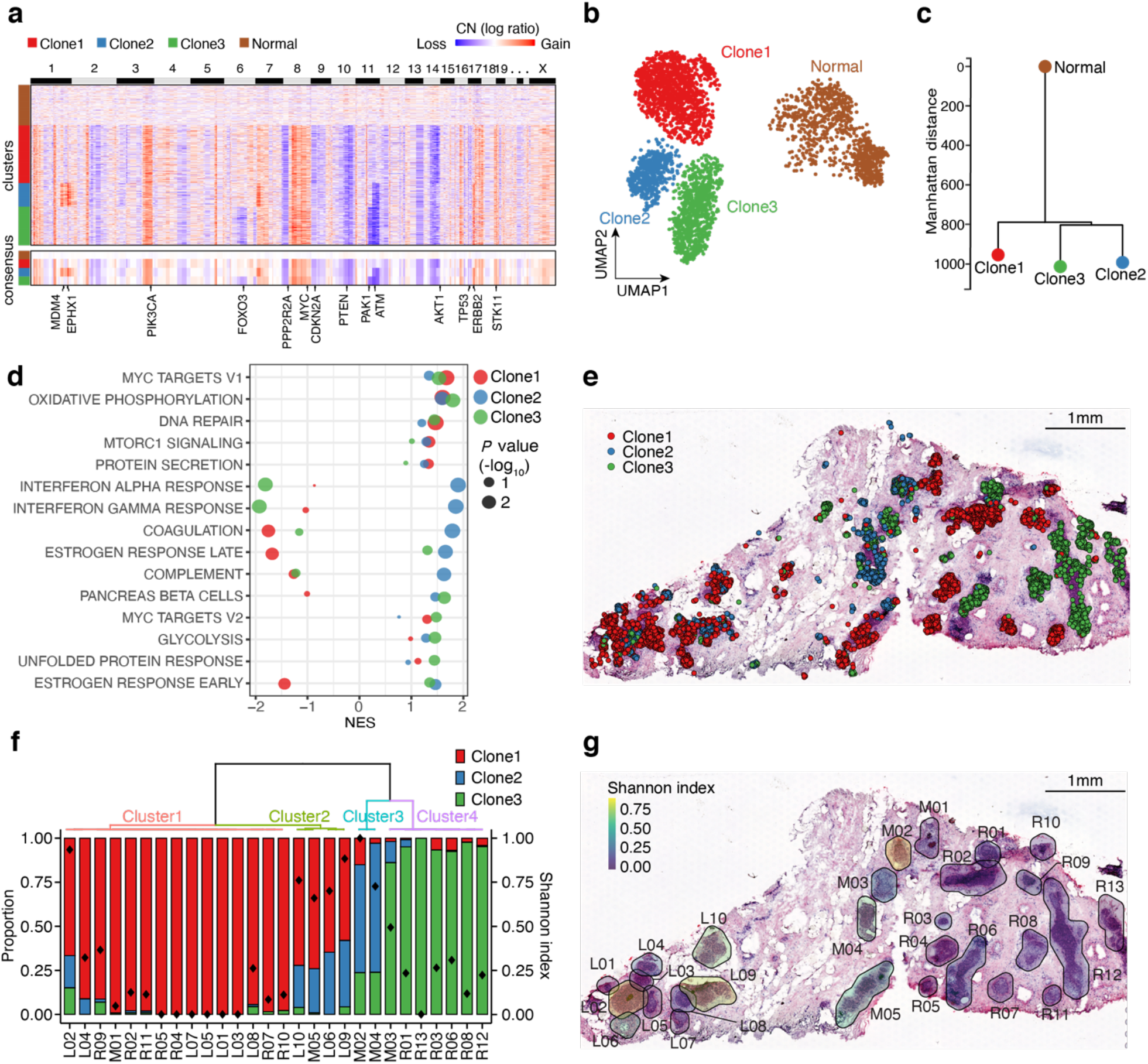
CellTrek identifies the spatial subclone heterogeneity in DCIS1. **a**, A heatmap of copy number (CN) profiles inferred by CopyKAT on the scRNA-seq data in DCIS1. The lower part represents a consensus CN profile of each cluster with some breast cancer-related genes annotated. **b**, CN-based UMAP of DCIS1. **c**, Phylogenetic tree based on the consensus CN profiles. **d**, Hallmark GSEA analysis of the expression data from three tumor subclones. **e**, Spatial cell charting of three tumor subclones using CellTrek. **f**, Tumor subclonal compositions within different ducts. The diamond symbol in each bar represents the Shannon index which measures the diversity of tumor subclones. **g**, H&E image of the DCIS tissue section with Shannon diversity index for each duct.

To understand the spatial distribution of three tumor subclones, we applied CellTrek to the scRNA-seq and ST data. Most of the tumor cells mapped to the DCIS regions on the H&E slide (Fig. 4e, g and Supplementary Data 3). Moreover, different tumor subclones mapped to different ductal regions, reflecting extensive spatial intratumor heterogeneity^44^. Specifically, clone2 was localized mostly to the middle (M) ducts, while clone3 was located primarily on the right (R) ducts and clone1 was spread across many ductal regions (Fig. 4e, g). Unsupervised clustering of the ST tumor spots identified five ST clusters that showed spatial and gene expression concordance to the tumor CellTrek map (Supplementary Fig. 7c-e). Based on the subclonal compositions of each duct, we performed a clustering analysis and calculated Shannon index, resulting in four major ductal clusters with different subclone compositions and spatial patterns (Fig. 4f). Overall, ducts from the right part of the tissue displayed low clonal diversities, while some ducts from the middle and left regions showed higher clonal diversities (Fig. 4g).

We further investigated the spatial co-expression patterns of the tumor cells using SCoexp and identified three gene modules (K1, K2 and K3). The K1 module was high in Clone1 and enriched with actin-related pathways (Supplementary Fig. 8a, e, f). CellTrek displayed that cells with high K1 scores corresponded to tumor clone1 spatially (Supplementary Fig. 8g). By contrast, K2 was high in Clone2 and Clone3 and was enriched with response to estradiol, mammary gland duct morphogenesis and some catabolic processes (Supplementary Fig. 8h-j). Interestingly, the K3 module was highly active in proliferating tumor cells and associated with cell cycle related processes (Supplementary Fig. 8b-d, k, l). The spatial mapping of the K3 score showed that proliferating tumor cells were mostly located near the peripheral regions of several ducts (Supplementary Fig. 8m). Taken together, these data show that the CellTrek toolkit can delineate topological maps of different tumor subclones and their expression programs in a DCIS tissue.

### Spatial tumor-immune microenvironment of a DCIS tissue

In another DCIS sample with synchronous invasive components (DCIS2), we profiled 3,748 single cells (10X Genomics) and 2,063 ST spots (Visium, 10X Genomics). Unsupervised clustering and DE analyses identified 10 clusters, including three epithelial clusters, endothelial, pericytes, fibroblasts, myeloid, NK/T, B and plasmacytoid dendritic cells (pDC) (Supplementary Fig. 9a, b). CopyKAT revealed an aneuploid epithelial cluster with CNAs (epithelial3) (Supplementary Fig. 9c). Histopathological analysis of the H&E image identified 11 ductal regions with tumor cells (T1-T11) and intervening areas that contained stromal and immune cells (Fig. 5a). To study the tumor-immune microenvironment, we focused on aneuploid cells and immune cells from the scRNA-seq data (Fig. 5b). Using CellTrek, we mapped most of the aneuploid cells to the histologically defined DCIS regions and immune cells to areas surrounding ducts and stromal regions (Fig. 5c and Supplementary Data 4). Interestingly, we found that some immune cells, including T, B, myeloid cells and pDC, were aggregated in the areas directly outside of the ducts, especially T1, T2, T6 and T7. Combining the CellTrek result with the H&E image, we posited the existence of tertiary lymphoid structures (TLS) in these regions. To further investigate this question, we calculated ST spot-level TLS scores^45, 46^ and found that spots with high TLS scores often corresponded to the mixed immune cell aggregations in our CellTrek map (Fig. 5c,d). Furthermore, we found that the ST-level TLS scores positively correlated with the charted immune cell counts (Pearson’s R = 0.36, *P* = 1.2e-10) (Fig. 5e). Together, these results show that CellTrek is capable of reconstructing the spatial tumor-immune microenvironment based on the scRNA-seq and ST data.

**Fig. 5.**
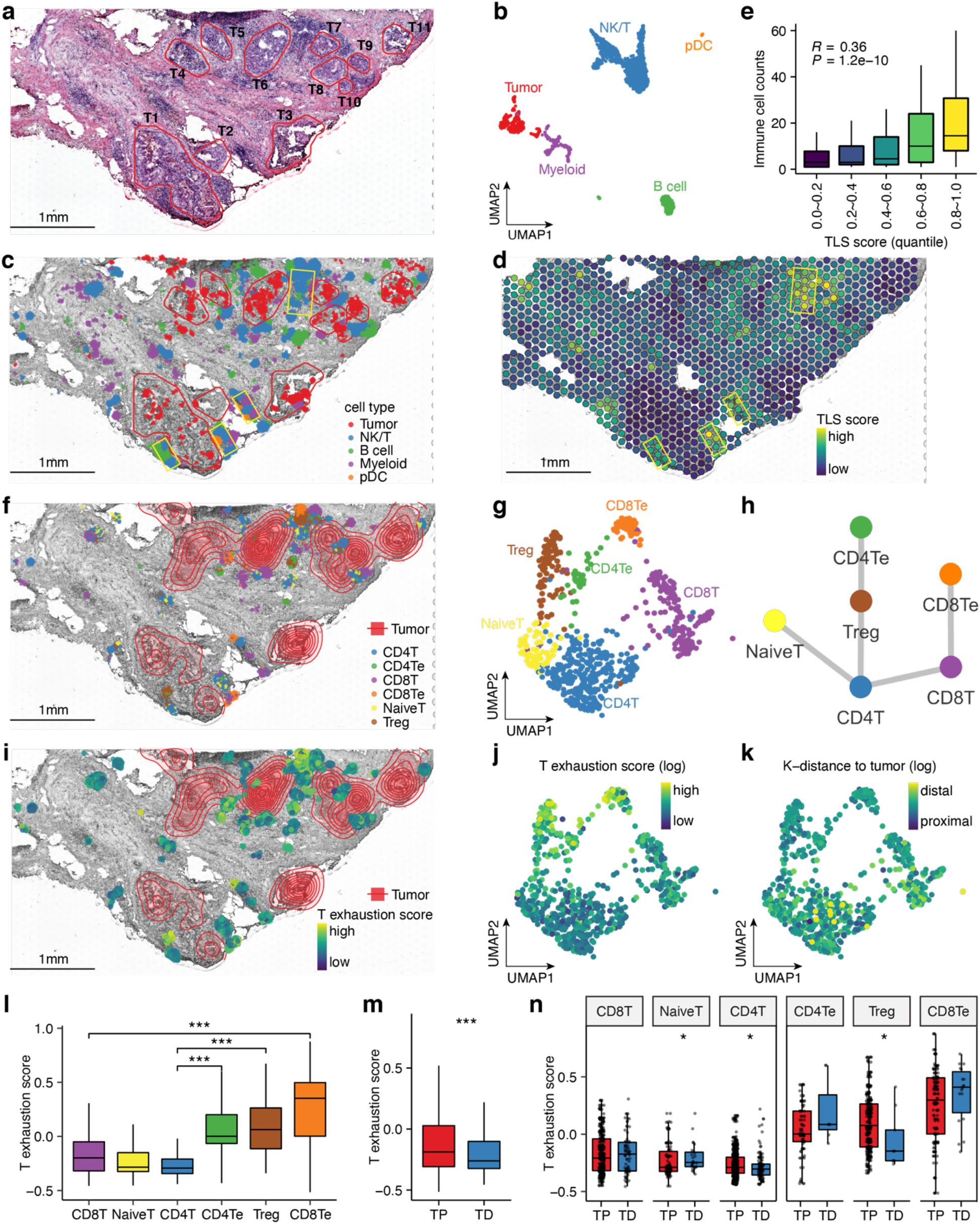
CellTrek displays the spatial tumor-immune microenvironment in DCIS2. **a**, H&E image of the tissue section from the DCIS2 patient. Histopathological annotations of tumor regions are highlighted in red circles with labels from T1 to T11. **b**, UMAP of DCIS2 scRNA-seq data (tumor cells, B cells, NK/T cells, myeloid and pDC cells). **c**, CellTrek spatial mapping of tumor cells, B cells, NK/T cells, myeloid and pDC cells. Yellow boxes highlight potential locations of tertiary lymphoid structures (TLS) with aggregation of mixed immune cells. **d**, ST spot-level TLS signature scores. **e**, Boxplot showing the association between CellTrek-based immune cell counts and ST spot TLS score quantiles. **f**, CellTrek spatial mapping of different T cell states. The contour plot represents the tumor cell densities. **g**, UMAP of scRNA-seq data showing different T cell states. **h**, Spatial colocalization graph of T cell states using SColoc. **i**, CellTrek spatial mapping of the T exhaustion scores. **j**, UMAP of T cells showing the exhaustion scores. **k**, UMAP of T cells showing the spatial K-distances to their 15 nearest tumor cells. **l**, Boxplot comparing the T cell exhaustion scores between different T cell states. **m**, Boxplot comparing the T cell exhaustion scores between T cells proximal to tumor cells (TP) and T cells distal to tumor cells (TD). **n**, Boxplot comparing the T cell exhaustion scores between TP and TD within each T cell state. In **l, m** and **n**, * indicates *P* < 0.05, *** indicates *P* < 0.01, *** indicates *P* < 0.001 using Wilcoxon rank-sum test. Boxplots show the median with interquartile ranges (25–75%); whiskers extend to 1.5X the interquartile range from the box.

Next, we found that some T cells were proximal to and some were distal to the tumor regions. We further analyzed the T cells and re-clustered them into six cell states, including the Naive T (NaiveT), CD4^+^ T (CD4T), CD8^+^ T (CD8T), regulatory T cells (Treg), exhausted CD4^+^ T (CD4Te) and exhausted CD8^+^ T (CD8Te) (Fig. 5g and Supplementary Fig. 9d). We investigated the distribution of these T cell states in the CellTrek map. Notably, the Tregs, CD4Te and CD8Te cells were mostly proximal to the tumor cells (Fig. 5f). We further constructed a spatial graph within the T cells and found that cells from the same lineages tended to colocalize spatially (Fig. 5h). We calculated T exhaustion scores and found that T cells with high exhaustion scores tended to localize near the tumor areas (Fig. 5i). K-distances of the T cells to their 15 nearest tumor cells showed an opposite trend to the T exhaustion scores on the UMAP (Fig. 5j, k). As expected, the immunosuppressive T cells (Treg, CD4Te and CD8Te) had higher exhaustion scores compared to the non-suppressive T cells (Fig. 5l). We binarized T cells to tumor distal (TD) and tumor proximal (TP) groups based on their K-distances and found that the TP group showed significantly higher exhaustion scores than the TD group (*P* = 1.1e-4, Fig. 5m), suggesting the presence of immunosuppressive microenvironment near the DCIS ductal regions. We also found a similar trend in which TP had higher exhaustion scores compared to TD for the CD4T and Treg cells and an opposite trend for the NaiveT cells (Fig. 5n). Importantly, the TD groups contained only few immunosuppressive T cells, which is consistent with our finding that exhausted T cells tend to colocalize near DCIS regions (Fig. 5n).

Re-clustering of the myeloid cells identified four cell states, including conventional dendritic cells (cDCs), monocytes and two macrophage subpopulations (Macro1 and Macro2; Supplementary Figs. 9e and 10a). CellTrek projected most of the cDCs to the tumor proximal areas (Supplementary Fig. 10a). The spatial graph showed that the Macro2 cells were colocalized with Macro1 and cDC (Supplementary Fig. 10b). We then calculated the K-distance of myeloid cells to the tumor cells (Supplementary Fig. 10c) and found that the cDCs displayed the lowest K-distances overall, while the Macro1 cells had higher K-distances. The K-distance density plot showed a similar trend (Supplementary Fig. 10d). We further examined the spatial co-expression of the Macro1 cells and identified two major gene modules (K1, K2) and one minor module using SCoexp. The K1 module was more active in macrophages from tumor distal regions (Supplementary Fig. 10e) and correlated with multiple *C1Q* genes, *HAVCR2, CD74, HLA-DRA*, etc. (Supplementary Fig. 10g). Conversely, the K2 module showed an opposite spatial pattern (Supplementary Fig. 10f) and correlated with *CHIT1, CSTB, APOC1, MARCO* and others (Supplementary Fig. 10g).

To orthogonally validate the spatial distribution of tumor and immune cells inferred by CellTrek, we performed immunofluorescence (RNAscope) experiments with targeted probes for tissue slides from DCIS2 and another DCIS sample (DCIS3). This data showed that the DCIS tumor cell areas had high expression of *ERBB2*, while *TAGLN* marked the basal epithelial layers of the ducts (Supplementary Fig. 11a,b). Furthermore, immune suppressive T cell markers, including *CTLA4* and *FOXP3*, had high expression near the DCIS areas in DCIS2 (Supplementary Fig. 11b,c) which is consistent with the CellTrek results. Similarly, in DCIS3, we found immunosuppressive T cells with *CTLA4* and *FOXP3* near the ducts (Supplementary Fig. 11d-f). Additionally, this data showed that B cells (*MS4A1)*, monocytes/macrophages (*CD68*) and dendritic cells (*CD1C*) were also near the DCIS ductal regions, suggesting the presence of TLS (Supplementary Fig. 11g), and was consistent with the CellTrek results for DCIS2. In contrast, fewer immune cells were observed in the normal lobular epithelial areas in the same tissue section, particularly for the immune suppressive T cell markers (Supplementary Fig. 11h,i). These data confirmed our findings on the DCIS tumor-immune microenvironment that were inferred using CellTrek.

## Discussion

Here, we report a novel computational tool, CellTrek, for reconstructing a spatial cellular map based on scRNA-seq and ST data. In contrast to conventional deconvolution approaches^11-14^, CellTrek provides a new paradigm that directly projects single cells to their spatial coordinates in tissue sections and therefore takes full advantage of the scRNA-seq data. We also developed two downstream computational modules (SColoc and SCoexp) to further analyze the CellTrek results. By reconstructing a cellular spatial map, CellTrek provides several advantages. First, it provides a flexible way to investigate any feature of individual cells (e.g., cell types/states, pseudotime) in a spatial manner, while most ST deconvolution approaches can only decompose spots into cell types, and cannot achieve single cell level feature mapping. Second, CellTrek is very flexible and can take any cell-location probability/similarity matrix as an input to reconstruct a cellular map, thus enabling further downstream analyses. Third, by utilizing a metric learning approach and a non-linear interpolation, CellTrek allows more accurate cell charting in a higher spatial resolution. Finally, with the development of higher spatial-resolution sequencing technologies, CellTrek is fully capable of charting single cells to other spatial sequencing data to provide even higher spatial granularity.

We first benchmarked the CellTrek performance using the simulated and *in situ* datasets and then evaluated the accuracy and robustness under different data conditions. By applying the CellTrek toolkit to two “well-established” datasets from mouse brain and kidney, we demonstrated its capability in recovering the topological structures of different cell types. We further showed that CellTrek can identify high-resolution substructures by mapping categorical (i.e., cell states) and continuous features (i.e., pseudotime) to the tissue sections. SColoc can also reconstruct the spatial relationships of different cell types into a graph, which can be further leveraged for cell-cell communication analysis. Moreover, SCoexp can detect spatial co-expression modules within multiple cell types, showing topological patterns in the tissue sections.

In our study we performed matched scRNA-seq and ST experiments of two DCIS samples and applied the CellTrek toolkit to delineate the spatial distribution of the tumor subclones in different ductal regions and the topological organization of the tumor-immune microenvironment. In DCIS1, we found that three tumor subclones were localized to different ducts with different levels of clonal diversity. Although morphological and genomic intratumor heterogeneity have previously been observed, here we report spatial heterogeneity within the ductal network in the DCIS tissue^44, 47-49^. In DCIS2, CellTrek accurately mapped tumor and immune cells, and indicated the presence of TLS enriched with immune cells near the DCIS regions. Further analyses of T cells and myeloid cells revealed their spatial localization relative to the tumor cells. These findings were orthogonally validated using RNAscope.

While CellTrek is a powerful tool for analyzing scRNA-seq and ST data, it has several notable limitations. First, CellTrek can have sparse cell mapping in some tissue areas as we showed in the simulation data. To overcome this problem, one can 1) collect tissues with higher cellular density for ST analysis; 2) sequence more cells or integrate multiple scRNA-seq datasets. Second, CellTrek maps cells to their most similar spots based on a sparse graph, which requires ST spots with relative high cell purities. A simulation of increasing spatial randomness (decreasing ST spots purities) showed that CellTrek could potentially over-simplify the spatial complexity for “less-organized” tissue structures. Finally, there is a risk of over-interpreting the data only based on CellTrek since it is a computational inference tool. Although relatively stringent parameters are used as a default to control for false positives, orthogonal validation is recommended to confirm biological findings.

In the future, CellTrek could be improved by including image recognition or deep learning approaches for cell segmentation and identification. Additionally, epigenetic regulation is of great interest in developmental biology and cancer research. Therefore, another future direction is to adapt CellTrek for epigenome data (e.g., scATAC-seq) to understand spatial epigenetic regulation in the tissue sections. Overall, we expect that CellTrek will have a multitude of applications for studying basic biology and human disease in spatial context, as applying scRNA-seq and ST experiments to the same tissues is becoming ever more commonplace.

## Supporting information

Supplementary Bundle

## Data and software availability

The scRNA-seq and ST data were submitted to the Gene Expression Omnibus (GEO): GSE181254. The CellTrek software toolkit is available at GitHub: https://github.com/navinlabcode/CellTrek.

## Acknowledgements

This work was supported by grants to N.E.N. from the NIH National Cancer Institute (RO1CA240526, RO1CA236864), the CPRIT Single Cell Genomics Center (RP180684) and the Chan-Zuckerberg Initiative (CZI) SEED Network Grant (CZF2019-002432) and the PRECISION Cancer Grand Challenge Grant. N.E.N. is an AAAS Fellow, AAAS Wachtel Scholar, Damon-Runyon Rachleff Innovator, Andrew Sabin Fellow, and Jack & Beverly Randall Innovator. This study was supported by the MD Anderson Sequencing Core Facility Grant (CA016672). The work was also supported by a Damon-Runyon Quantitative Biology Postdoctoral Fellow to R.W. We thank Jingye Wang, Xiaoling He and Julian Wei for their unwavering support.

## Author Contributions

R.W. and S.H. developed the CellTrek method, analyzed data prepared figures and wrote the manuscript. S.B. and E.S. performed single cell and ST experiments. M.H. processed data. A.T. and S.K. collected the DCIS tissue samples, interpreted data and managed the IRB protocols. K.C. provided input on the CellTrek method and manuscript. N.N. managed the project and wrote the manuscript.

## Methods

### CellTrek toolkit

#### CellTrek

Using ST and scRNA-seq data, CellTrek first uses a reference-based co-embedding approach from the Seurat package^25^ with scRNA-seq data as the reference and the ST data as the query. From the co-embedded data, CellTrek then trains a multivariate random forest (RF) model with a default of 1,000 trees on ST data using *rfsrc* from the randomForestSRC package^50^ with the following formula:

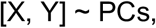

where X and Y are spatial coordinates of ST spots, and PCs are the top principal components (default = 30). Additionally, CellTrek introduces a non-linear 2D interpolation approach from the akima package to augment the ST spots. The trained RF model is applied to the ST-scRNA co-embedding data to produce an RF-based distance matrix. RF distance is calculated by

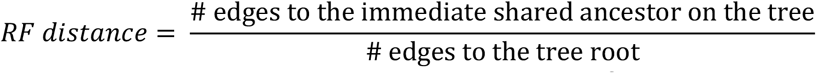

This distance metric provides a semi-parametric measurement of the similarities between data points in their feature space while supervised by the spatial coordinates. Based on the RF distance matrix, CellTrek further constructs a sparse graph by filtering distances larger than a threshold and matching the closest cell-spot pairs using mutual nearest neighbors. This sparse graph enables a flexible cell charting scheme with certain degrees of redundancy considering that one ST spot contains multiple cells and different ST spots could consist of similar cells. In addition to the default machine learning-based distance matrix, CellTrek is designed to be compatible with any cell-location similarity/probability matrix computed from other methods as an input, such as novoSpaRc^21^, thus extending the compatibility and scalability of CellTrek. Using the sparse spot-cell graph, CellTrek assigns the coordinates from ST spots to their connected neighboring cells. To avoid the cell clumping problem, CellTrek applies point repulsion algorithm using *circleRepelLayout* from the packcircles package. Additionally, we also developed an interactive plotting function (*celltrek_vis*) for the CellTrek cell map visualization which allows mapping any continuous or categorical cell features to the spatial map with different colors and shapes provided. This plotting function also allows interactive manual selection and annotation of cells.

#### Scoloc

To recapitulate the colocalization of different cell types on the CellTrek results, we developed the SColoc module that provides three different approaches to calculate cell closeness, i.e., KL-divergence (KL), Delaunay triangulation (DT) and K-nearest neighbor distance (KD) considering different tissue structures or study goals. For the KL-based approach, SColoc calculates a 2D grid kernel density for each cell type using *kde2d* from the MASS package with default *h* equals to spatial distance between two neighbor ST spots and n = 25. Then, SColoc calculates the KL-divergence on the 2D density between each pair of two cell types. KL-based SColoc works well for detecting the global closeness of large spatial structures, for example, the neuron layer structure in the brain. For DT-based approach, it first builds a 2D Delaunay triangulation network using *delaunayn* from the geometry package based on the CellTrek result. To confine the network complexity, SColoc can further filter edges with distances larger than a certain threshold. Then, on the cell type level, SColoc calculates neighboring cell counts on the DT network. Between any pair of cell types, SColoc calculates a log odds ratio that represents the colocalization of these two cell types. This approach shows better performances in capturing connections between cell types when local cellular structures are more of interest (for example, DCIS samples). To further simplify the connections, SColoc will also build a minimum spanning tree (MST). The above calculation procedures are performed repetitively on bootstrap samples (default 20 iterations). Both bootstrap closeness matrix and MST consensus matrix are produced for graph visualization. SColoc also provides an interactive visualization function (*scoloc_vis*) to render a graph representation of cell colocalization using either MST consensus or bootstrap closeness as input with flexible tuning thresholds to simplify the complexity.

#### Scoexp

To identify potential spatial-relevant gene co-expression modules, especially within cell types of interest, we developed the SCoexp module based on the CellTrek results. For cell of interests, SCoexp first calculates a spatial distance matrix between each cell and converts it into a spatial kernel matrix *W* using radial basis function (RBF) with a default sigma equals to the distance between two neighbored ST spots. Using this spatial kernel matrix *W* and a cell gene expression matrix *X*, SCoexp calculates a spatial-weighted gene cross-correlation matrix based on the following formula:

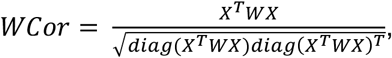

where *diag()* is the diagonal vector of a matrix. Based on the spatial-weighted gene cross-correlation matrix, SCoexp uses two co-expression module detection approaches, i.e., consensus clustering (CC)^22^ and weighted correlation network analysis (WGCNA)^23^. CC approach applies the ConsensusClusterPlus package with a default K through 2 to 8 and default repetition of 20. Then for the identified gene clusters, a within-cluster filtering step removes low consensus and low correlation genes in each module. WGCNA approach identifies co-expression modules with the normalized *WCor* matrix as an input. Similarly, a within-module filtering can be applied to remove low correlation genes. For the identified gene modules, we applied the Seurat *AddModuleScore* to calculate a cell-level module activity score which can be investigated on the CellTrek spatial map.

### Mouse data acquisition and analysis

#### Mouse brain data

For the mouse brain scRNA-seq (Smart-seq2)^18^ and ST data (10X Genomics Visium), we downloaded the Seurat objects from https://satijalab.org/seurat/articles/spatial_vignette.html. For the scRNA-seq data, we randomly subsampled 8,000 cells from the original dataset and performed a standard data analysis procedure including log normalization, scaling, variable genes selection (n = 5,000) using *vst*, dimension reduction using PCA and UMAP. On each cell type, we then applied *dbscan* (v 1.1-5)^51^ with minPts = 20 and eps = 0.5 to filter out outliers in the UMAP. For the ST data, we performed similar analysis processing (log-normalization, scaling, variable genes identification and dimensionality reduction) as the scRNA-seq data.

#### Mouse hippocampus data

The mouse hippocampus scRNA-seq data^31^ was downloaded from the Seurat website (https://satijalab.org/seurat/articles/spatial_vignette.html#slide-seq-1). We subsampled 20,000 cells from the data and performed *SCTransform*. Mouse hippocampus Slide-seq data^30^ was installed through the Seurat package. We subsampled 10,000 spots and performed *SCTransform*. For the Slide-seq data, we then conducted dimensionality reduction using PCA, UMAP and clustering analysis using Seurat.

#### Mouse kidney data

For the mouse kidney 3’ scRNA-seq data (10X Genomics), we downloaded the filtered gene count matrices from GEO (GSE129798), and the cell type annotation from the original paper^32^. The data included single cells from three different dissection zones, i.e. cortex, outer medulla and inner medulla. We processed the scRNA-seq data following the similar procedures as the mouse brain data. We also subsampled 8,000 cells and filtered out outlier cells using group-wise *dbscan*. On proximal tubule and distal tubule cells, we conducted trajectory analysis using the Monocle2 package (v 2.14.0)^27^ with raw count data as an input and the negative binomial distribution as the expressionFamily argument. Highly variable genes with mean expression more than 0.25 were used to perform gene ordering and dimensionality reduction based on *DDRTree* function. A principal graph was generated and cells were ordered along the pseudotime trajectory. The mouse kidney ST data was downloaded from the 10X Genomics website (https://www.10xgenomics.com/resources/datasets/) and analyzed by the same procedure as described for the mouse brain ST process workflow.

### DCIS tissue collection and sequencing

#### ScRNA-seq experiments

The two DCIS samples were obtained from the MD Anderson Cancer Center. The study was approved by the Institutional Review Board (IRB) and tissue was procured with informed consent from the patients. The tumors were stained with hematoxylin and eosin (H&E) and evaluated by pathology, in which DCIS1 was classified as a pure DCIS sample and DCIS2 as a synchronous DCIS-IDC tissue. The estrogen receptor (ER) and progesterone receptor (PR) status of the samples were determined by immunohistochemistry (IHC), which showed that DCIS1 was ER and PR positive, while DCIS2 was ER positive and PR negative. We used our previous protocol to prepare viable single cell suspensions^42^. Briefly, fresh tissue samples were dissociated into viable single cell suspensions by enzymatic dissociation using collagenase A and trypsin. The fresh tissues were also embedded into Optimal Cutting Temperature (OCT) and snap-frozen. The viable cell suspensions were used as input material for scRNA-seq using the Single-Cell Chromium 3′ protocol by V2 (10X Genomics CG00052, PN-120237) and V3 (10X Genomics CG000183, PN-1000075) chemistry reagents. The final libraries containing barcoded single-cell transcriptomes were sequenced at 100 cycles using the S2 flowcell on the Novoseq 6000 system (Illumina).

#### ST Visium experiments

Fresh tissues from the two DCIS patients were cut to proper size and embedded in cryomold (Fisher #NC9542860) by OCT compound (Fisher #1437365) on dry ice and stored in -80°C in sealed bags. Frozen OCT embedded DCIS cryosections were cut to 12μm in the cryostat (Thermo Scientific Cryostar NX70) with specimen head temperature at -17 °C and blade temperature at -15 °C. The cut sections were placed within a capture area of the Visium spatial slide (10X Genomics PN-1000184). The slide was permeabilized for 12 minutes according to the Visium Spatial Tissue Optimization protocol (10X Genomics CG000238). Imaging of the stained slides was performed on the Nikon Eclipse Ti2 system. Finally, the ST libraries were constructed by following the Visium Spatial Gene Expression protocol (10X Genomics CG000239) and sequenced at 200 cycles by S1 flowcell on the Novoseq 6000 system (Illumina).

#### RNAscope experiments

RNAscope probes were ordered from Advanced Cell Diagnostics (ACD) for the following genes: *ERBB2, ACTG2, TAGLN, CTLA4, FOXP3, CD3D, CD4, CD8A, MA4A1, CD1C* and *CD68*. Cryosections of two snap-frozen DCIS samples (OCT embedded) were cut at a thickness of 10um and used to performed RNA in situ hybridization assay with the RNAscope Multiplex Fluorescent v2 kit according to the manufacturer’s instructions (Cat# 323110 and 323120) with following modifications: tissue sections were fixed in 10% NBF for 1hr at 4°C, all washing steps were increased to 3 times (3-5min each wash), areas enclosed by hydrophobic barrier were 0.75”x0.75”, for all reagent steps 150ul were dispensed, tissues were treated with Protease III for 15min at RT, Opal dyes were used at 1/750-1/2250 (Akoya Biosciences FP1487001KT, FP1488001KT, FP1495001KT FP1497001KTP), kit DAPI stain was replaced by a 1/2000 working stock in PBS (Invitrogen D1306 in DMF, 5mg/ml), and slides were mounted in Prolong diamond (Invitrogen #P36970). Fluorescent Images were scanned using a motorized stage on the Nikon Eclipse Ti2 microscope with 20X objective and analyzed with the Nikon NIS-Elements AR software (5.30.04).

### DCIS scRNA-seq data analysis

FASTQ files were first preprocessed using the *Cell Ranger* 3.1.0 pipeline (10X Genomics) with default arguments and mapped to the *GRCh38* reference genome to construct count matrices. Unique molecular identifier (UMI) counts were then processed using the Seurat package (v 3.2)^25^. For the two DCIS samples, cells with less than 700 unique feature counts were filtered. We also filtered cells that had the percentage of mitochondrial counts more than 15%. Counts were then normalized using the *NormalizeData* with default *LogNormalize* method. Afterwards, normalized counts were scaled and centered using *ScaleData* function. 2,000 variable genes were found using *FindVariableFeatures* function and principal component analysis (PCA) was conducted using *RunPCA* with default parameters. *ElbowPlot* was used to determine the number of PCs for the downstream analyses. 10 neighbor cells and the top 20 PCs were used for neighbor finding using *FindNeighbors*. Cell clusters were identified using *Louvain* algorithm of *FindCluster* function with the top 20 PCs and a resolution of 0.6. We ran *RunUMAP* to visualize cell manifold in a 2D space. Differentially expressed (DE) genes were identified using *FindAllMarkers*. Cell identities were annotated based on a combination of two strategies: 1) Top markers for each cluster based on the DE gene analysis; 2) Canonical cell type specific marker expression using *FeaturePlot*. We used dbscan with minPts = 5 and eps = 0.5 to remove outliers for each cell type based on the UMAP. Then, we utilized CopyKAT (v17) to infer the copy number for each cell^42^. For DCIS1, the inferred copy number profiles were plotted using ComplexHeatmap (v 2.2.0)^52^ and used to generate copy number alterations (CNA)-based UMAP using the uwot package (v 0.1.8) on the Manhattan distance matrix. We then applied dbscan to identify tumor subclones in a CNA-based UMAP space. Consensus copy number profiles for the tumor subclones and normal cells were then used to construct a phylogenetic tree based on neighbor joining tree in the ape package (v 5.4)^53^ and the normal cell was chosen as root. Gene set enrichment analysis was performed on three tumor subclones using the fgsea package^43^. Cell cycle scores for cells were calculated using the *CellCycleScore* module from Seurat and assigned as in G1, G2/M or S phase. For DCIS2, to further define NK/T and myeloid cell subtypes, these cells were extracted and re-analyzed using the similar workflow. DE analysis was performed and cell subtypes were annotated based on the top gene expression and canonical maker expression. To characterize T cell exhaustion, we used an exhaustion gene signature including *PDCD1, CTLA4, LAG3, HAVCR2, CD244, CD160* and *TIGIT*.

### DCIS ST data analysis

Sequencing data were first preprocessed with *Space Ranger* v1.0.0 and mapped to the *GRCh38* reference genome. Similar to the scRNA-seq data analysis workflow, the ST data were subsequently processed using Seurat. We filtered out spots with UMI counts less than 100. The UMI counts were normalized using *NormalizeData* function with *LogNormalize followed by ScaleData*. 2,000 variable genes were found using *FindVariableFeatures* function and PCA was performed using *RunPCA*. For DCIS1, to study tumor spots, we selected the ST spots covering the ductal areas based on histopathology and conducted an unsupervised clustering using the top 20 PCs at resolution of 0.5. For DCIS2, we cropped out the damaged areas of the original tissue slide which resulted in a total of 965 spots left. The downstream analyses were based on the cropped tissue section. For ST spots, the TLS score was calculated based on a 12-gene signature^46^ using *AddModuleScore* function in Seurat.

### Benchmarking and simulations

#### Simulation data

We used R package Splatter^54^ to generate the simulation scRNA-seq data with 6,000 cells, 5,000 genes and 5 cell groups. We set lib.loc = 8.5, lin.scale = 0.4 and dropout.type = ‘group’. We used the function *splatSimulatePaths* to simulate a sequential manifold with path.skew = 0.1, de.facLoc = 0.1 and de.facScale = 0.8 for all groups. We then generated a customized spatial area and assigned cells to the locations using optimal transport between the top 2 principal components and spatial locations using the transport package^55^. ST spots were generated using a 30 by 20 grid and spots outside the spatial area were then dropped. Each ST spot aggregated the gene expression from its 5 nearest cells. A final ST data with 394 spots was used.

#### Drosophila embryo data

This data is from the Berkeley *Drosophila* Transcription Network Project (BDTNP) (https://www.fruitfly.org/)^21^. Based on the FISH data of *Drosophila* embryo, they generated 3,309 cells with the expression level of 84 landmark genes. We filtered 5 outlier cells and applied the Seurat analysis pipeline to identify 7 cell groups at resolution = 0.25. Similarly, we generated the ST data with a final of 483 spots.

#### Mouse embryo data

We downloaded the seqFISH data of mouse embryo 1 from https://marionilab.cruk.cam.ac.uk/SpatialMouseAtlas/^24^. We downsampled 6,000 cells and removed the low-quality group and groups with less than 50 cells. We finally used a single cell data of 5,852 cells and 351 genes for test. ST data was correspondingly generated with a final of 779 spots. We sub-selected a group of foregut cells and ran Monocle2 for trajectory analysis.

#### Different conditions of simulation

Based on the original scRNA-seq simulation data, we further simulated different data conditions to test the robustness of CellTrek. In simulation-1, we simulated different read counts for both scRNA-seq and ST data by dropping a certain number of reads from a proportion of cells and spots. The detailed dropout parameters were described in Supplementary Table 1. This simulation will not only produce data with different library size and read depth but also introduce batch effects between scRNA-seq and ST data since the dropouts were conducted independently on these two data sets. In simulation-2, we simulated spatial randomness by swapping cells in a spatially manner. Specifically, we first added a Gaussian noise with a customized deviation to the original cell coordinates and then assigned the cells with spatial noise back to the original tissue structure (Supplementary Table 2, Supplementary Fig. 4c). The increasing Gaussian noise deviation will increase the spatial randomness compared to the raw reference structure (Supplementary Fig. 4d, left). Corresponding ST data was then generated at each condition. This simulation also produced a decrease of cell type purities within ST spots (Supplementary Fig. 4d, right). In simulation-3, we simulated different tissue densities by spatially down-sampling cells. We first set-up some spatial marker points (Supplementary Table 3 and Supplementary Fig. 4g), and then calculated cell distances to the marker points and converted into RBF kernel densities. We then down-sampled cells based on their kernel densities. This simulation can generate different spatial sparseness in different areas. ST data was then generated correspondingly. To quantitatively evaluate the CellTrek performance, we compared the CellTrek results to their spatial cellular reference under different conditions using cell type-level KL-divergences and correlation analysis on spatial coordinates. Permutation tests of 100 times were performed to generate null distributions.

### CellTrek and downstream analyses

#### Simulation and benchmarking data

For the simulated scRNA-seq and ST data, we first ran *traint* to co-embed the data into a shared feature space with default parameters. We tested the RF-distance distribution between ST and scRNA-seq data and determined a threshold around 1% quantile which is 0.35 (350 out of 1,000). Using this threshold, we ran *celltrek* on the co-embedded *traint* data with following parameters: intp_pnt = 2,000 spots, nPCs = 20, ntree = 1,000, top_spot = 4, spot_n = 5 and repel_r = 10 with 10 iterations. For NVSP-CellTrek, we first ran *novoSpaRc* using reference-guided mode using the same default parameters following the protocol from https://github.com/rajewsky-lab/novosparc/blob/master/reconstruct_drosophila_embryo_tutorial.ipynb. On the *gw* matrix from *novoSpaRc*, we ran a negative log transformation and determined a threshold of 13 (around 1% quantile of the distance). We then ran *celltrek_from_dist* with following parameters: top_spot = 4, spot_n = 5, dist_cut = 13, and reprel_r = 20. For SrtCT, we ran *FindTransferAnchors* from Seurat with ST data as the reference and scRNA-seq data as the query and reduction of “cca”. We then applied *TransferData* to transfer the ST coordinates from ST data to single cells with default parameters. For *Drosophila* and mouse embryo data, similar to the simulation data, we performed CellTrek, NVSP-CellTrek and SrtCT to analyze the single cell and generated ST data using similar parameter settings. For CellTrek simulation analysis of different data conditions, we first filtered low quality ST and scRNA-seq data using nFeature > 10. We set intp = FALSE due to the running time and repel_r was set to 20 considering no spatial interpolation here. All parameters were fixed across different simulations and conditions.

#### Mouse brain

We subset the frontal cortex region in the ST data for single cell spatial charting. For the normalized ST and scRNA-seq data, we first ran *traint* to co-embed the data into a shared feature space with default parameters. We tested the RF-distance distribution between ST and scRNA-seq data and determined a threshold close to 1% quantile which is 0.5 (500 out of 1,000). Using this threshold, we ran *celltrek* on the co-embedded *traint* data with following parameters: intp_pnt = 10,000 spots, nPCs = 20, ntree = 1,000, top_spot = 5, spot_n = 10 and repel_r = 20 with 10 iterations. For NVSP-CellTrek, we ran *novoSpaRc* using reference-guided mode. Similarly, we ran a negative log transformation on the *gw* matrix and determined a threshold of 14 by testing the distribution (1% quantile). We then ran *celltrek_from_dist* with following parameters: top_spot = 5, spot_n = 10, dist_cut = 14, and reprel_r = 20. SrtCT was performed with the same parameters as the simulation study. To evaluate different cell charting approaches, we benchmarked SrtLT as our reference. Specifically, we used *SelectIntegrationFeatures* to select 2,000 features between ST and scRNA-seq data. Then we used *FindTransferAnchors* with scRNA-seq data as the reference, ST data as the query data and reduction of “cca”. Then we applied *TransferData* to transfer the cell type labels from scRNA-seq data to the ST spots. For comparison, for each cell type, we converted all charting results (CellTrek, NVSP-CellTrek and SrtCT) to a spatial grid density with h = 10, n = 50 and coordinates limits fixed to the range of ST spots coordinates. For SrtLT, we first applied a customized function that generates pseudo-points based on probabilities and used the same approach to calculate a spatial grid density for each cell type. We then focused on cell types that had 1) more than 10 spots with label transfer probability > 0.5 and 2) more than 20 cells in the scRNA-seq data. For each cell type, we used total normalization and calculated the KL-divergence between any cell chart density and the SrtLT reference density. To calculate the cell spatial colocalization, we applied *scoloc* on the CellTrek results using “KL” approach with cell_min = 15, eps = 1e-50 and boot_n = 20. We calculated the mean of bootstrap KL matrices and converted it into a similarity matrix. This matrix was then used for heatmap with *ward*.*D2* clustering. For the graph visualization, we used the MST consensus matrix as input while setting a consensus cutoff at 0.3. For the glutamatergic neurons, we calculated spatial K-distance by setting the L2/3 IT cell type as the reference and k = 10. K-distance was then normalized by the maximum values. For L5 IT cells, we performed the spatial co-expression analysis. We first filtered mitochondria, ribosomal, and highly sparse (non-zero proportion less than 20%) genes and calculated highly variable genes which resulting in a total of 2,000 genes. We then used a consensus clustering based spatial co-expression analysis using *scoexp* with following parameters: sigm = 140, avg_con_min = 0.5, avg_cor_min = 0.4, zero_cutoff = 3, min_gen = 50, max_gen = 400 and maxK = 8. After gene modules were identified, we applied *AddModuleScore* from Seurat to calculate the cell module activity score with nbin = 10 and ctrl = 20. We performed Gene Ontolgy (GO) enrichment analysis for the two modules using the clusterProfiler package^56^. To identify genes correlated with the co-expression modules, we used a customized *FindCorMarkers* function in CellTrek based on the Spearman correlation.

#### Mouse hippocampus

For the Slide-seq and scRNA-seq on mouse hippocampus, *traint* and *celltrek* were consequentially conducted with norm = “SCT”, nPCs = 50, dist_thresh = 0.65, top_spot = 5, spot_n = 2, and repel_r = 10.

#### Mouse kidney

Similar to the mouse brain data, we performed three different cell charting approaches for comparison, i.e., CellTrek, NVSP-CellTrek, and SrtCT to analyze this scRNA-seq and ST data. For CellTrek, the dist_thresh was set to 0.6 considering the distance distribution. For NVSP-CellTrek, we selected a distance cutoff of 13.5. We also benchmarked SrtLT and compared these cell charting results using KL-divergence on the spatial grid density at h = 20 and n = 40. For the spatial colocalization analysis, we used *scoloc* with “DT” approach on the CellTrek results. The diagonal of the mean matrix was replaced with NA to emphasize only the colocalization between different cell types. The MST consensus matrix was employed for graph visualization at a cutoff of 0.2. Similarly, the similarity matrix was used for heatmap with the *ward*.*D2* clustering. We applied *scoexp* with avg_cor_min = 0.3 on the distal tubule cells to identify the spatial co-expression modules. We also performed GO enrichment analysis for these gene modules. To evaluate cellular interactions between different cell types of mouse kidney, we applied CellChat (v 1.1.3)^37^ to infer ligand-receptor interactions from the scRNA-seq data. We used the normalized count data as an input and followed the CellChat tutorial with default parameters and CellChatDB.mouse as the interaction database. Cellular interactions were visualized using *netVisual_circle* function. Next, in order to obtain a more specific interaction results, we leveraged our spatial SColoc graph. We binarized the MST consensus matrix at 0.2 as a spatial weight matrix. We calculated the element-wise product of CellChat cellular interaction matrix and the weight matrix. Ligand-receptors interaction examples were then plotted using *netVisual_chord_gene* function.

#### DCIS1

We applied *traint* followed by *celltrek* with dist_thresh = 0.5, top_spot = 5, spot_n = 10, and repel_r = 5 to construct a spatial cell map. To calculate the diversities of different ducts, we assigned the tumor cells to their closest ducts with spatial distances less than 60. Then Shannon index was calculated using *entropy*.*empirical* from the entropy package and rescaled to 0∼1. To identify the similarity between the ST spots of the annotated ducts and the tumor subclones identified from the scRNA-seq data, we conducted DE analyses for the ST clusters and tumor subclones, respectively. Genes with adjusted *P* values (Bonferroni correction) less than 0.1 were selected to perform Pearson’s correlation between these two modalities on average log fold-change values. Spatial co-expression was conducted using the *scoexp* “cc” approach with sigm = 60, avg_cor_min = 0.4, zero_cutoff = 3, min_gen = 50 and max_gen = 400. Similarly, GO enrichment analysis was also performed for three spatial gene modules.

#### DCIS2

We used the same CellTrek procedure as described for DCIS1. To test the association between ST-TLS scores and CellTrek immune cell numbers (i.e., NK/T, B, Myeloid and pDC), we first assigned the immune cells to their closest ST spots and counted the cell numbers. Next, we calculated *Spearman* correlation between spot-level TLS scores and the corresponding immune cell counts. Within the T cell group, we applied *scoloc* with “DT” approach to summarize cell colocalization. MST consensus was used for graph visualization. We calculated K-distance of T cells to the tumor cells with K = 15. The K-distance of the T cells were then binarized to TD and TP groups based on a Gaussian mixture model^57^. For myeloid cells, we applied the same analysis procedure as T cells for the cell colocalization and K-distance analysis. For macrophages, we also conducted a spatial co-expression analysis using *scoexp* and identified highly associated genes using *FindCorMarkers*.

