## Supplementary Bundle for "Spatial charting of single cell transcriptomes in tissues"

#### Wei and He et al.

#### Supplementary Materials

##### CONTENTS

**Supplementary Figure 1**
Overview of two downstream computational modules of the CellTrek toolkit…….Page 3

**Supplementary Figure 2**
Three spatial single-cell datasets for benchmarking…….………………..……….Page 4

**Supplementary Figure 3**
Benchmarking on three datasets with known spatial references…………………...Page 5

**Supplementary Figure 4**
Simulation study of CellTrek under different data conditions…………..………...Page 7

**Supplementary Figure 5**
Mouse brain and kidney co-expression module enrichment analysis and mouse kidney spatial constrained cell-cell interaction analysis…………………………..…..…...Page 9

**Supplementary Figure 6**
CellTrek analysis of mouse hippocampus Slide-seq data……………...…..…..….Page 11

**Supplementary Figure 7**
ScRNA-seq and ST analyses of DCIS1…………………………….……..…..…..Page 12

**Supplementary Figure 8**
Spatial co-expression patterns of tumor cells in DCIS1………..…………..…..…..Page 13

**Supplementary Figure 9**
ScRNA-seq data analysis of DCIS2……………………………………..…...….Page 14

**Supplementary Figure 10**
CellTrek analyses of myeloid cell states in DCIS2…………………...…..…...….Page 15

**Supplementary Figure 11**

Orthogonal validation of tumor and immune cell spatial distributions using RNAscope assay in DCIS2 and DCIS3……………………………..……………………..…..….Page 16

**Supplementary Table 1**
Parameter settings for different conditions in simulation-1…………..………..…..…..Page 18

**Supplementary Table 2**
Parameter settings for different conditions in simulation-2…………..………..…..…..Page 19

**Supplementary Table 3**
Parameter settings for different marker points in simulation-3…………..………..…..Page 20

**Supplementary Data 1**
Screenshot of the CellTrek result for mouse brain data…………..………..…..…..Page 21

**Supplementary Data 2**
Screenshot of the CellTrek result for mouse kidney data……..…..………..…..…..Page 22

**Supplementary Data 3**
Screenshot of the CellTrek result for DCIS1 data…………..……………………..…..Page 23

**Supplementary Data 4**
Screenshot of the CellTrek result for DCIS2 data…………..……………………..…..Page 24

#### Supplementary Figures

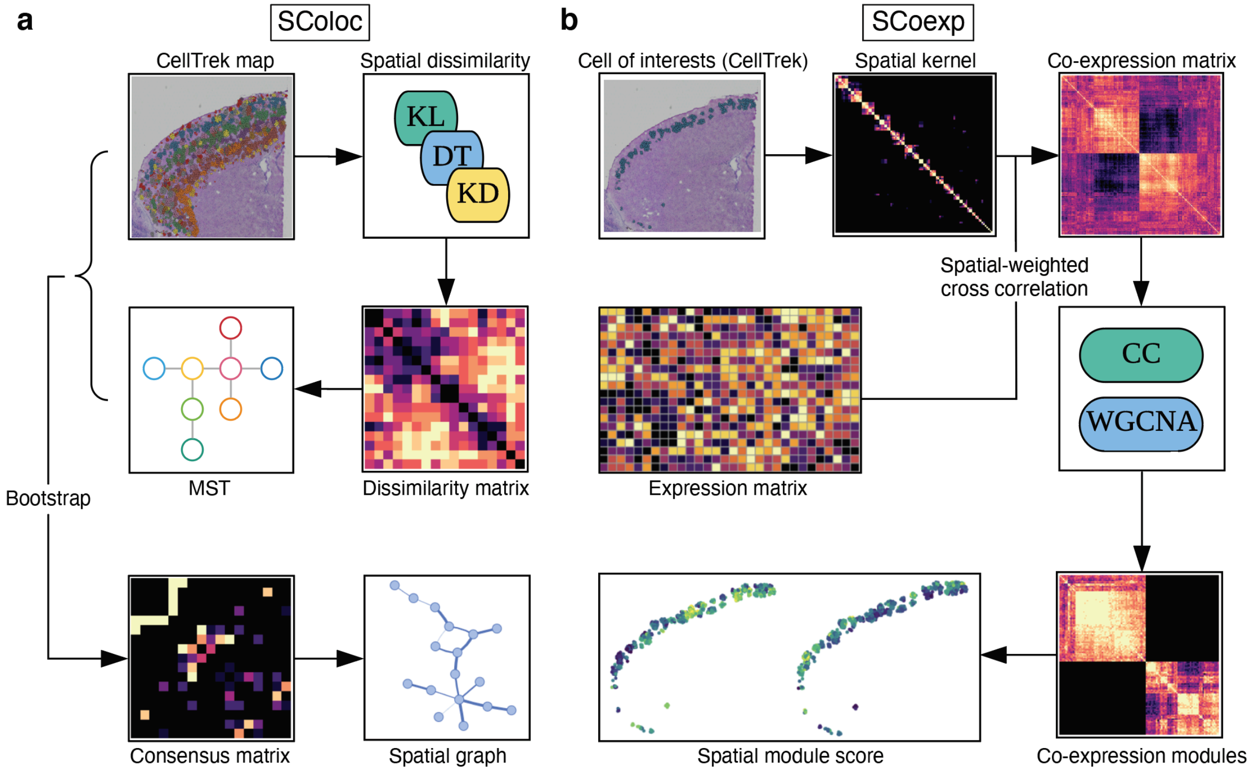


**Supplementary Figure 1 – Overview of two downstream computational modules of the CellTrek toolkit**

(**a**) The SColoc module. Based on the CellTrek result map, three different spatial dissimilarity methods, i.e., KL, DT, KD, can be applied to calculate a cell-type spatial dissimilarity matrix, and an MST is used to generate a tree structure. These steps are conducted repetitively on bootstrapped samples to calculate a consensus matrix on dissimilarity matrices or MSTs, which produces a final cell-type spatial graph representation. (**b**) The SCoexp module. For cells of interest, based on the CellTrek map, SCoexp first calculates a spatial kernel matrix using RBF based on their spatial distance. Next, based on the spatial kernel matrix and cell-gene expression matrix, SCoexp calculates the spatial weighted gene co-expression. Gene modules are then identified using CC or WGCNA. For the identified co-expression modules, module activity scores can be computed and mapped back to the CellTrek coordinates.

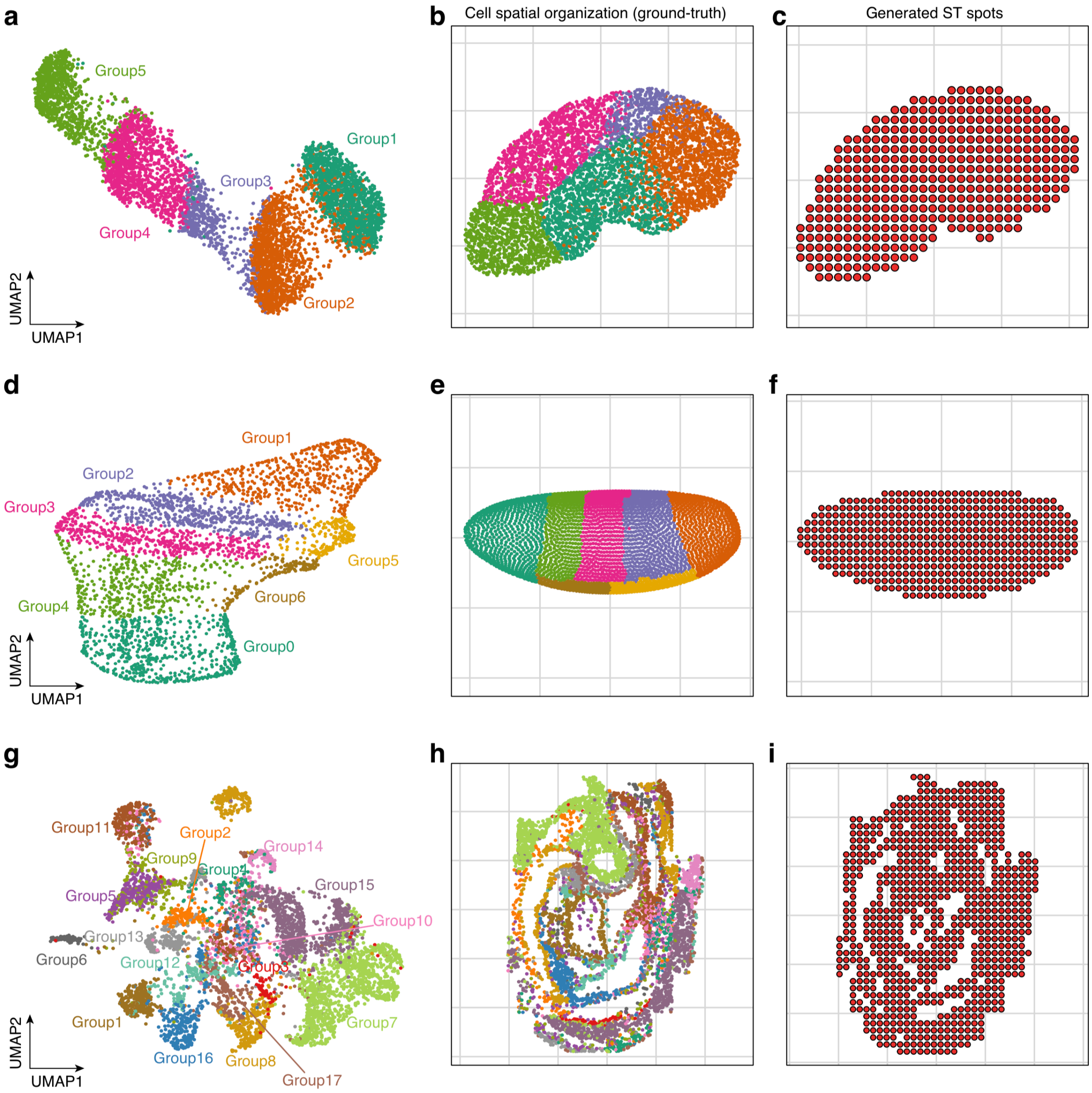


**Supplementary Figure 2 – Three spatial single-cell datasets for benchmarking**

(**a**) UMAP of a simulated scRNA-seq data with 5 cell groups. (**b**) Spatial organization of the simulated data as the ground truth. (**c**) ST data generated based on panel (**b**). (**d**) UMAP of *Drosophila* embryo FISH-generated single cell data. (**e**) Spatial organization of *Drosophila* embryo cells as the ground truth. (**f**) *Drosophila* embryo ST data generated based on panel (**e**). (**g**) UMAP of mouse embryo seqFISH data (Group1: Cardiomyocytes; Group2: Cranial mesoderm; Group3: Definitive endoderm; Group4: Dermomyotome; Group5: Endothelium; Group6: Erythroid; Group7: Forebrain midbrain hindbrain; Group8: Gut tube; Group9: Haematoendothelial progenitors; Group10: Intermediate mesoderm; Group11: Lateral plate mesoderm; Group12: Mixed mesenchymal mesoderm; Group13: Neural crest; Group14: Presomitic mesoderm; Group15: Spinal cord; Group16: Splanchnic mesoderm; Group17:Suface ectoderm). (**h**) Spatial organization of the mouse embryo cells as the ground truth. (**f**) mouse embryo ST data generated based on panel (**h**). In (**c**), (**f**) and (**i**), each ST spot aggregates the 5 nearest cells to generate the ST data.

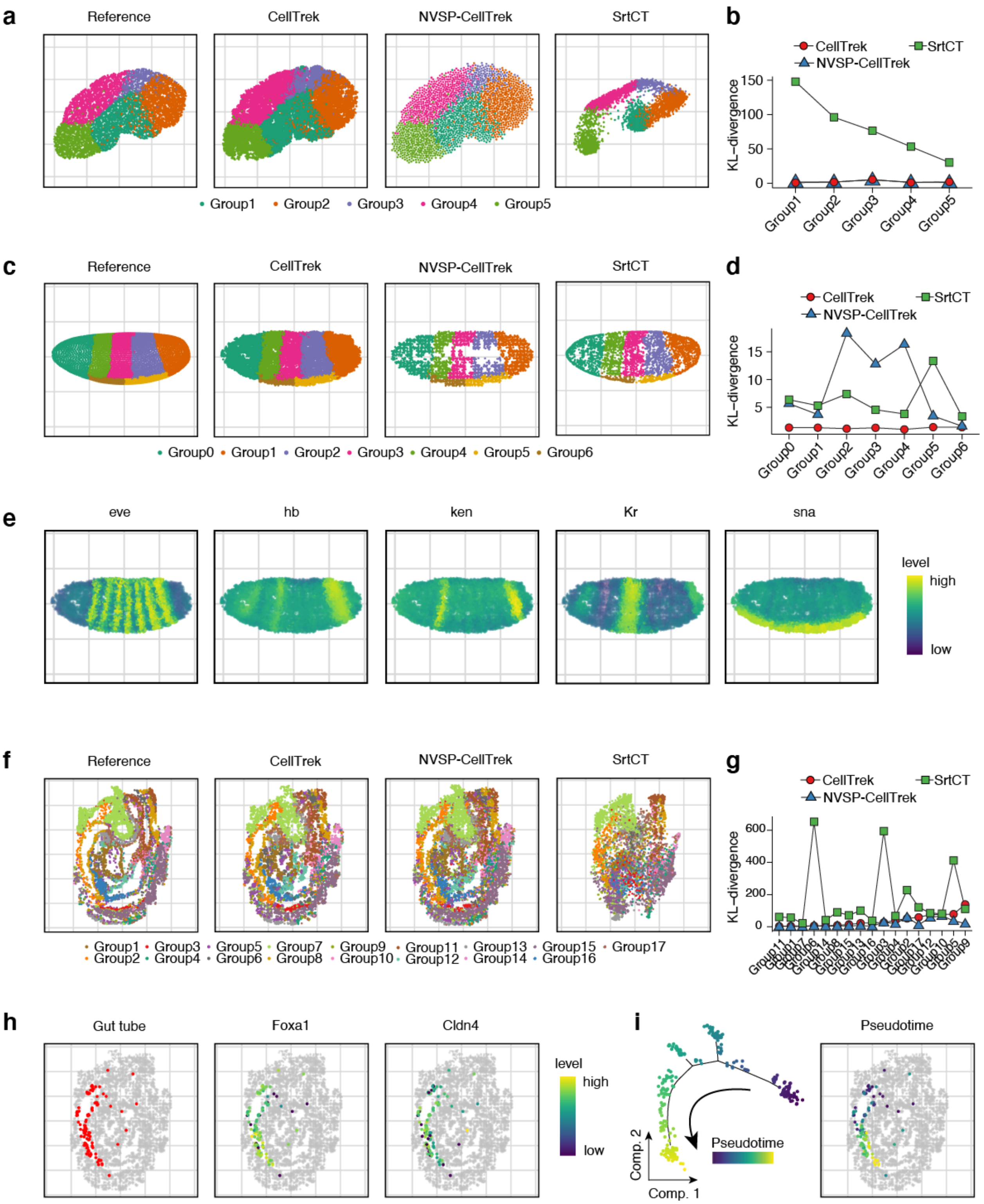
a
Reference
CellTrek
NVSP-CellTrek
SrtCT
Group1
Group2
Group3
Group4
Group5
b
CellTrek
SrtCT
NVSP-CellTrek
KL–divergence
c
d
Group0
Group6
e
eve
hb
ken
Kr
sna
level
high
low
f
g
h
Gut tube
Foxa1
Cldn4
i
Pseudotime
Comp. 2
Comp. 1

**Supplementary Figure 3 – Benchmarking on three datasets with known spatial references**

(**a**) Comparison of the ground-truth reference, CellTrek, NVSP-CellTrek and SrtCT results on the simulated data. (**b**) KL-divergence of spatial cell charting methods for each cell type compared with the ground-truth. (**c**) Comparison of the ground-truth reference, CellTrek, NVSP-CellTrek and SrtCT results on the *Drosophila* embryo data. (**d**) KL-divergence of spatial cell charting methods for each cell type compared with the ground-truth. (**e**) The expression of *Drosophola* embryogenic marker genes (*eve*, *hb*, *ken*, *Kr* and *sna*) on the CellTrek map. (**f**) Comparison of the ground-truth reference, CellTrek, NVSP-CellTrek and SrtCT results on the mouse embryo data (**g**) KL-divergence of three spatial cell charting methods for each cell type compared with the ground-truth. (**h**) The CellTrek map of a subgroup of gut tube (foregut cells) and the marker genes (*Foxa1* and *Cldn4*). (**i**) Trajectory analysis for foregut tube cells (left) and CellTrek spatial mapping of the pseudotime values in the tissue section (right).

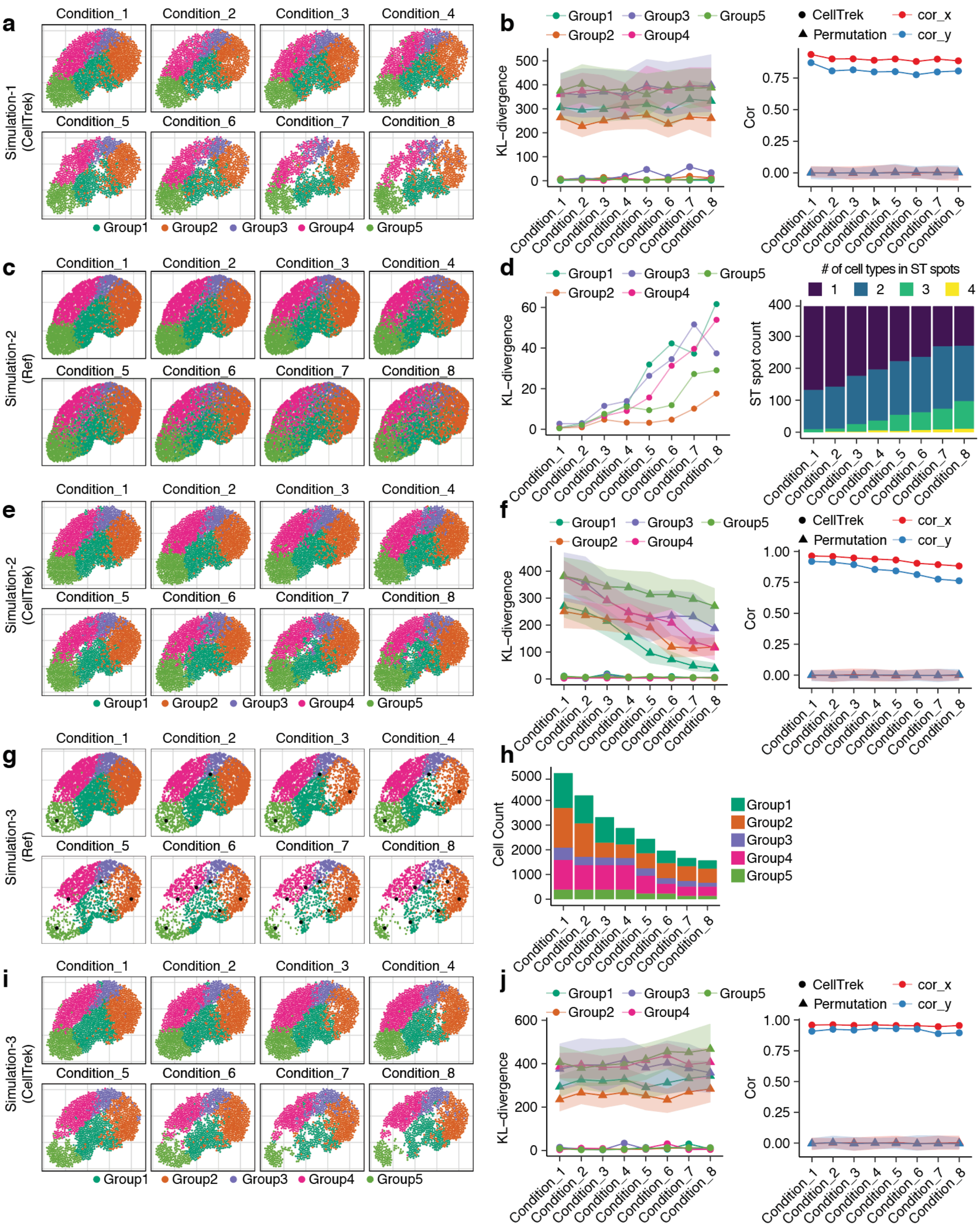
a
Simulation-1 (CellTrek)
Condition_1
Condition_2
Condition_3
Condition_4
Condition_5
Condition_6
Condition_7
Condition_8
Group1 Group2 Group3 Group4 Group5
b
KL−divergence
Cor
CellTrek
Permutation
cor_x
cor_y
c
Simulation-2 (Ref)
d
### of cell types in ST spots
1 2 3 4
ST spot count
e
Simulation-2 (CellTrek)
f
g
Simulation-3 (Ref)
h
Cell Count
i
Simulation-3 (CellTrek)
j

**Supplementary Figure 4 – Simulation study of CellTrek under different data conditions**

(**a**) CellTrek results under different gene counts in the simulated data. From condition1 to condition8, the gene counts decreased. (**b**) KL-divergence (left) and spatial coordinate correlation (right) of CellTrek and permutation test across different gene count conditions. (**c**) Simulation of the increasing spatial randomness of the reference data from condition1 to condition8. (**d**) KL-divergence of the simulated reference compared with the raw spatial cell organization (left) and the number of cell types in ST spot (right) across different spatial randomness conditions. (**e**) CellTrek results under different spatial randomness of the reference data. (**f**) KL-divergence (left) and spatial coordinate correlation (right) of CellTrek and permutation test across different spatial randomness conditions. (**g**) Simulation of the decreasing tissue spatial densities (cell numbers) of the reference data from condition1 to condition8. Black dots display the marker points for spatial down-sampling. (**h**) Bar plot showing the number of cells under different tissue spatial density conditions. (**i**) CellTrek results under different tissue densities of the reference data. (**j**) KL-divergence (left) and spatial coordinate correlation (right) of CellTrek and permutation test across different tissue spatial density conditions. Detailed parameter settings are provided in Supplementary Table 1-3. For the permutation tests, the ribbon shows the 95% confidence intervals.

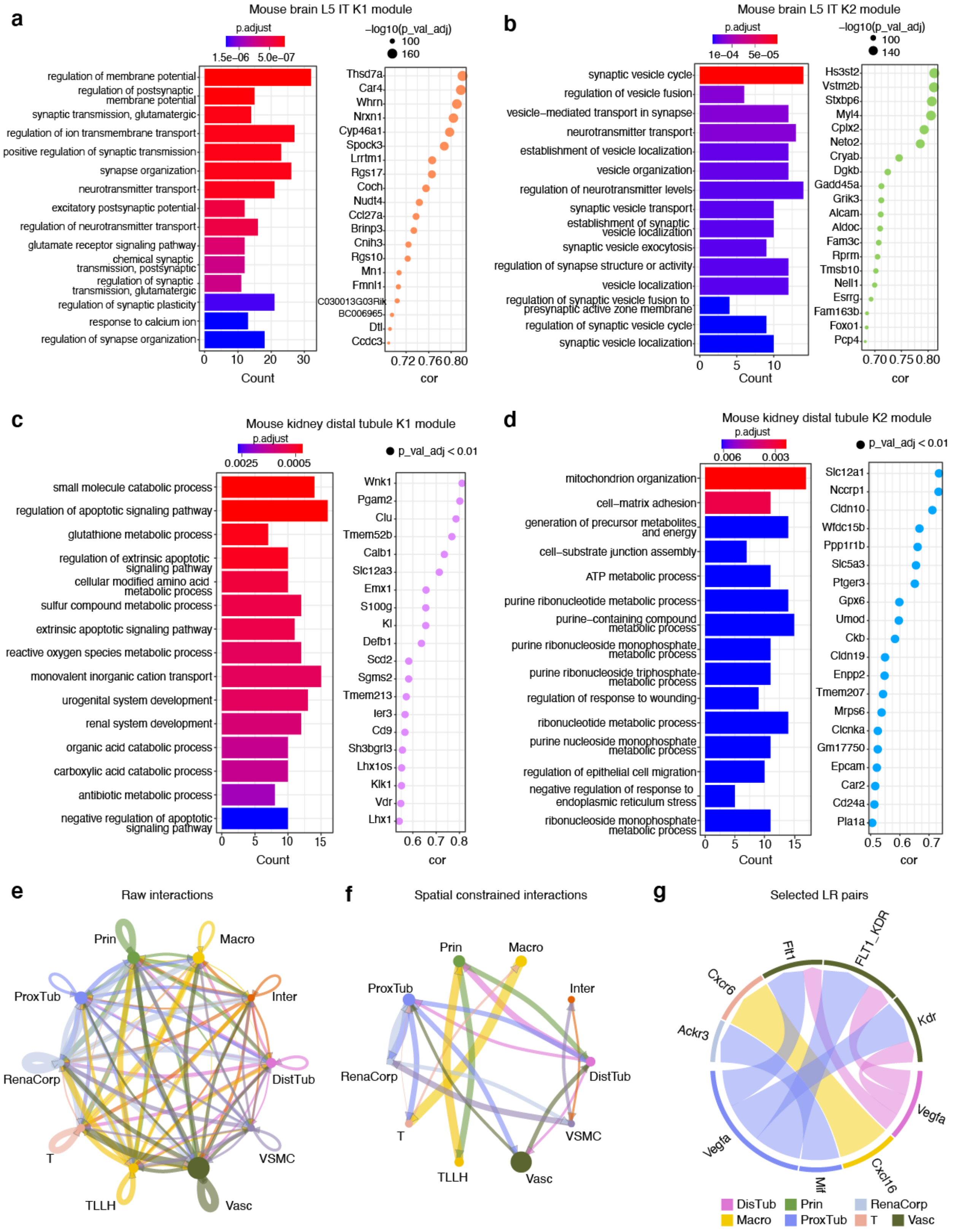
a
Mouse brain L5 IT K1 module
p.adjust
1.5e−06
5.0e−07
−log10(p_val_adj)
100
160
regulation of membrane potential
regulation of postsynaptic membrane potential
synaptic transmission, glutamatergic
regulation of ion transmembrane transport
positive regulation of synaptic transmission
synapse organization
neurotransmitter transport
excitatory postsynaptic potential
regulation of neurotransmitter transport
glutamate receptor signaling pathway
chemical synaptic transmission, postsynaptic
regulation of synaptic transmission, glutamatergic
regulation of synaptic plasticity
response to calcium ion
regulation of synapse organization
Count
Thsd7a
Car4
Whrn
Nrxn1
Cyp46a1
Spock3
Lrrtm1
Rgs17
Coch
Nudt4
Ccl27a
Brinp3
Cnih3
Rgs10
Mn1
Fmnl1
C030013G03Rik
BC006965
Dtl
Ccdc3
cor
b
Mouse brain L5 IT K2 module
p.adjust
1e−04
5e−05
−log10(p_val_adj)
100
140
synaptic vesicle cycle
regulation of vesicle fusion
vesicle−mediated transport in synapse
neurotransmitter transport
establishment of vesicle localization
vesicle organization
regulation of neurotransmitter levels
synaptic vesicle transport
establishment of synaptic vesicle localization
synaptic vesicle exocytosis
regulation of synapse structure or activity
vesicle localization
regulation of synaptic vesicle fusion to presynaptic active zone membrane
regulation of synaptic vesicle cycle
synaptic vesicle localization
Count
Hs3st2
Vstm2b
Stxbp6
Myl4
Cplx2
Neto2
Cryab
Dgkb
Gadd45a
Grik3
Alcam
Aldoc
Fam3c
Rprm
Tmsb10
Nell1
Esrrg
Fam163b
Foxo1
Pcp4
cor
c
Mouse kidney distal tubule K1 module
p.adjust
0.0025
0.0005
p_val_adj < 0.01
small molecule catabolic process
regulation of apoptotic signaling pathway
glutathione metabolic process
regulation of extrinsic apoptotic signaling pathway
cellular modified amino acid metabolic process
sulfur compound metabolic process
extrinsic apoptotic signaling pathway
reactive oxygen species metabolic process
monovalent inorganic cation transport
urogenital system development
renal system development
organic acid catabolic process
carboxylic acid catabolic process
antibiotic metabolic process
negative regulation of apoptotic signaling pathway
Count
Wnk1
Pgam2
Clu
Tmem52b
Calb1
Slc12a3
Emx1
S100g
Kl
Defb1
Scd2
Sgms2
Tmem213
Ier3
Cd9
Sh3bgrl3
Lhx1os
Klk1
Vdr
Lhx1
cor
d
Mouse kidney distal tubule K2 module
p.adjust
0.006
0.003
p_val_adj < 0.01
mitochondrion organization
cell−matrix adhesion
generation of precursor metabolites and energy
cell−substrate junction assembly
ATP metabolic process
purine ribonucleotide metabolic process
purine−containing compound metabolic process
purine ribonucleoside monophosphate metabolic process
purine ribonucleoside triphosphate metabolic process
regulation of response to wounding
ribonucleotide metabolic process
purine nucleoside monophosphate metabolic process
regulation of epithelial cell migration
negative regulation of response to endoplasmic reticulum stress
ribonucleoside monophosphate metabolic process
Count
Slc12a1
Nccrp1
Cldn10
Wfdc15b
Ppp1r1b
Slc5a3
Ptger3
Gpx6
Umod
Ckb
Cldn19
Enpp2
Tmem207
Mrps6
Clcnka
Gm17750
Epcam
Car2
Cd24a
Pla1a
cor
e
Raw interactions
Prin
Macro
ProxTub
Inter
RenaCorp
DistTub
T
VSMC
TLLH
Vasc
f
Spatial constrained interactions
Prin
Macro
ProxTub
Inter
RenaCorp
DistTub
T
VSMC
TLLH
Vasc
g
Selected LR pairs
Flt1
FLT1_KDR
Cxcr6
Kdr
Ackr3
Vegfa
Vegfa
Mif
Cxcl16
DisTub
Prin
RenaCorp
Macro
ProxTub
T
Vasc

**Supplementary Figure 5 – Mouse brain and kidney co-expression module enrichment analysis and mouse kidney spatial constrained cell-cell interaction analysis**

(**a-b**) GO enrichment analyses (left) and module-correlated genes (right) for mouse brain L5 IT K1 and K2 modules, respectively. (**c-d**) GO enrichment analyses (left) and module-correlated genes (right) for mouse kidney distal tubule K1 and K2 modules, respectively. (**e**) Summary of cell-cell interaction network of mouse kidney cells using CellChat. (**f**) Spatial constrained cell-cell interaction network of mouse kidney cells. (**g**) Chord plot showing some identified ligand-receptor pairs. The edge width in (**e**) and (**f**) represent the total interaction strength; and the arc width in (**g**) represent the interaction score calculated from CellChat. The direction of the arrows goes from the ligands to the receptors.

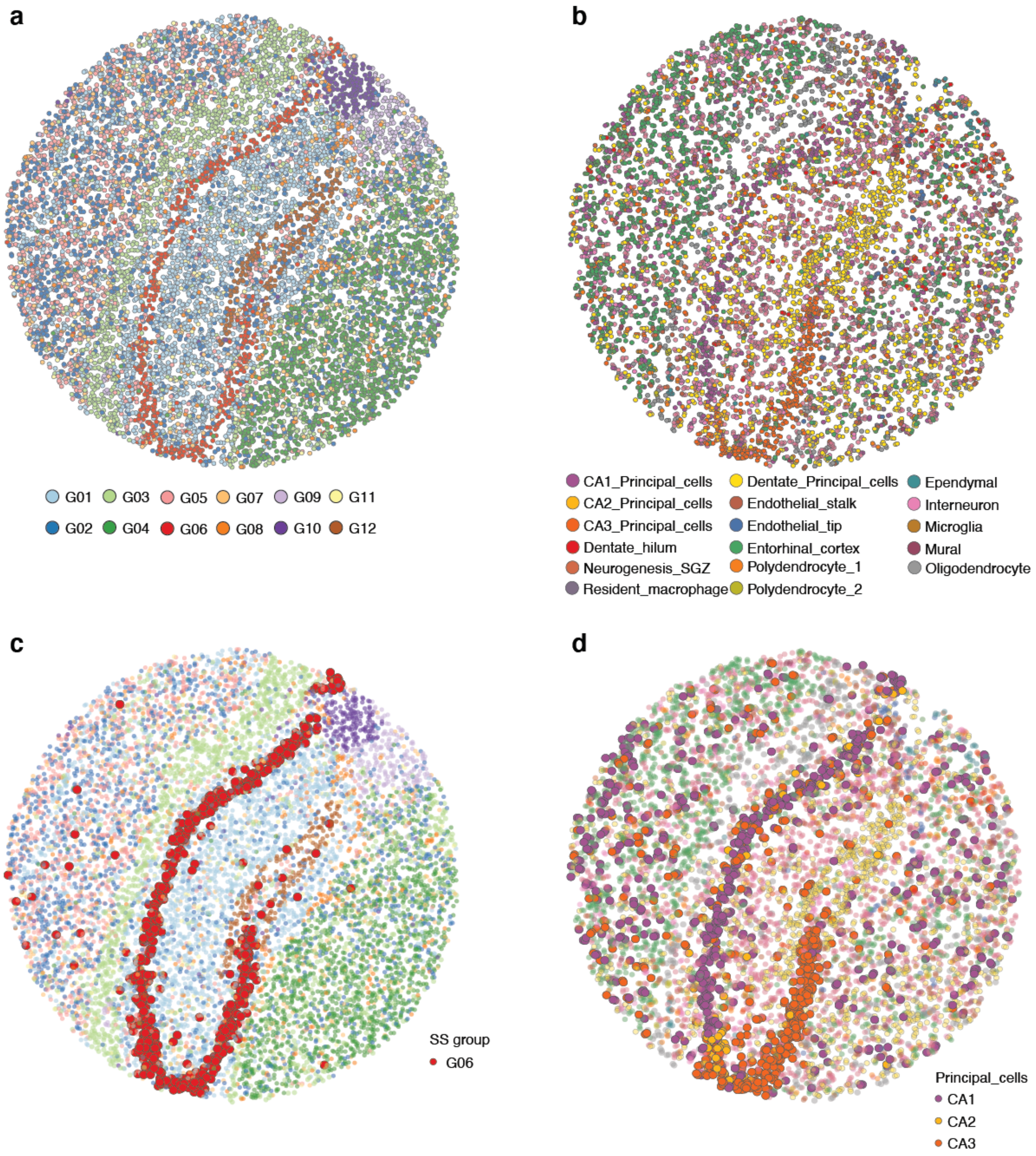


**Supplementary Figure 6 – CellTrek analysis of mouse hippocampus Slide-seq data**
(**a**) Unsupervised clustering of mouse hippocampus Slide-seq data identified 12 clusters. (**b**) CellTrek spatial mapping of scRNA-seq data to the tissue section. (**c**) Slide-seq data with G06 group highlighted. (**d**) CellTrek map with three principal cells (CA1, CA2 and CA3) highlighted which corresponded to G06.

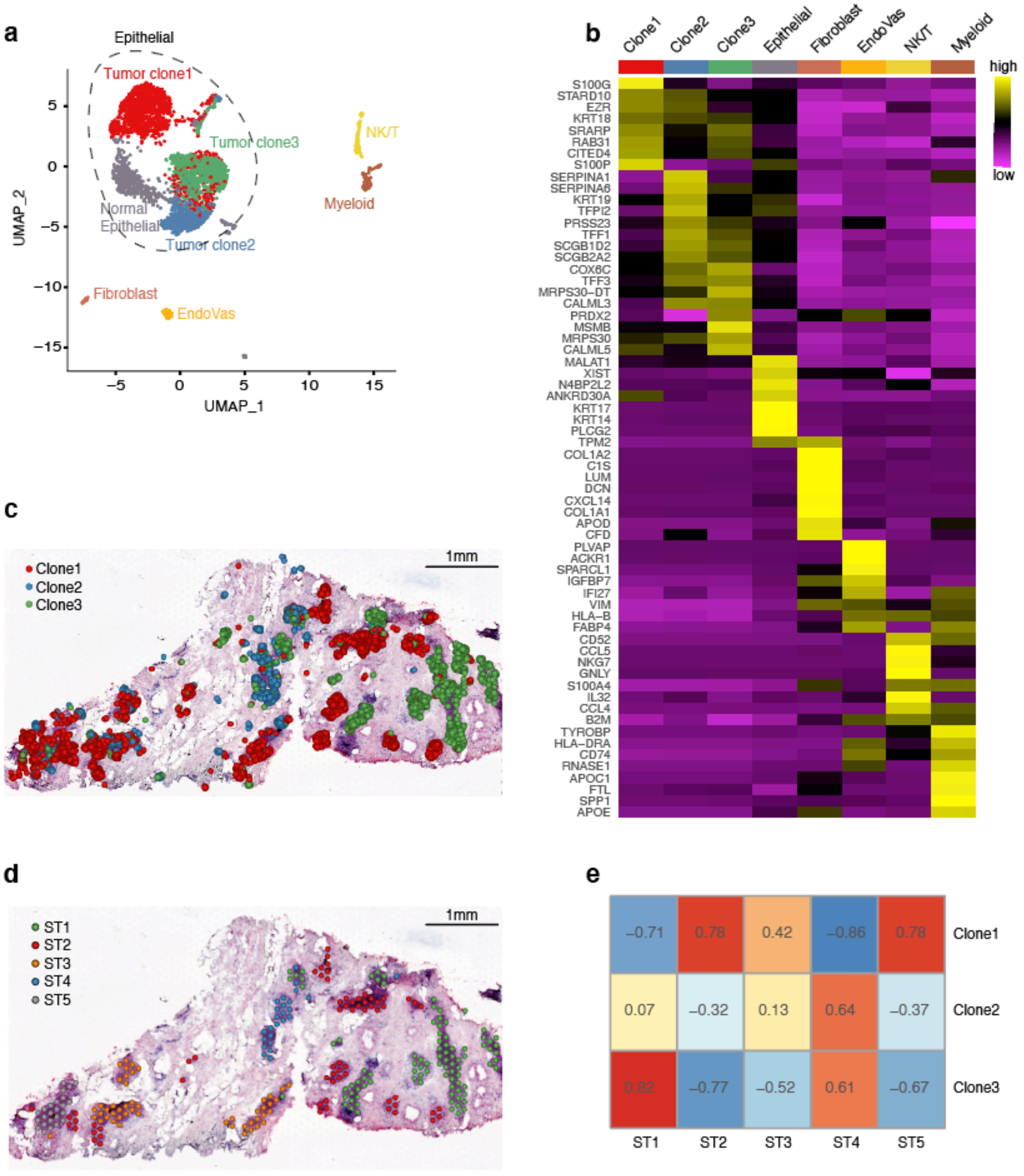


**Supplementary Figure 7 – ScRNA-seq and ST analyses of DCIS1**

(**a**) UMAP of DCIS1 scRNA-seq data. (**b**) Consensus heatmap of scRNA-seq data for top differentially expressed genes for each cell type. (**c**) CellTrek result of three tumor subclones (clone1-3) which is also presented in **Fig. 4e**. (**d**) Unsupervised clustering of ST tumor spots identified five clusters (ST1-5). (**e**) Gene expression-based Spearman correlation analysis between three tumor subclones in the scRNA-seq data (clone1-3) and five ST clusters (ST1-5).

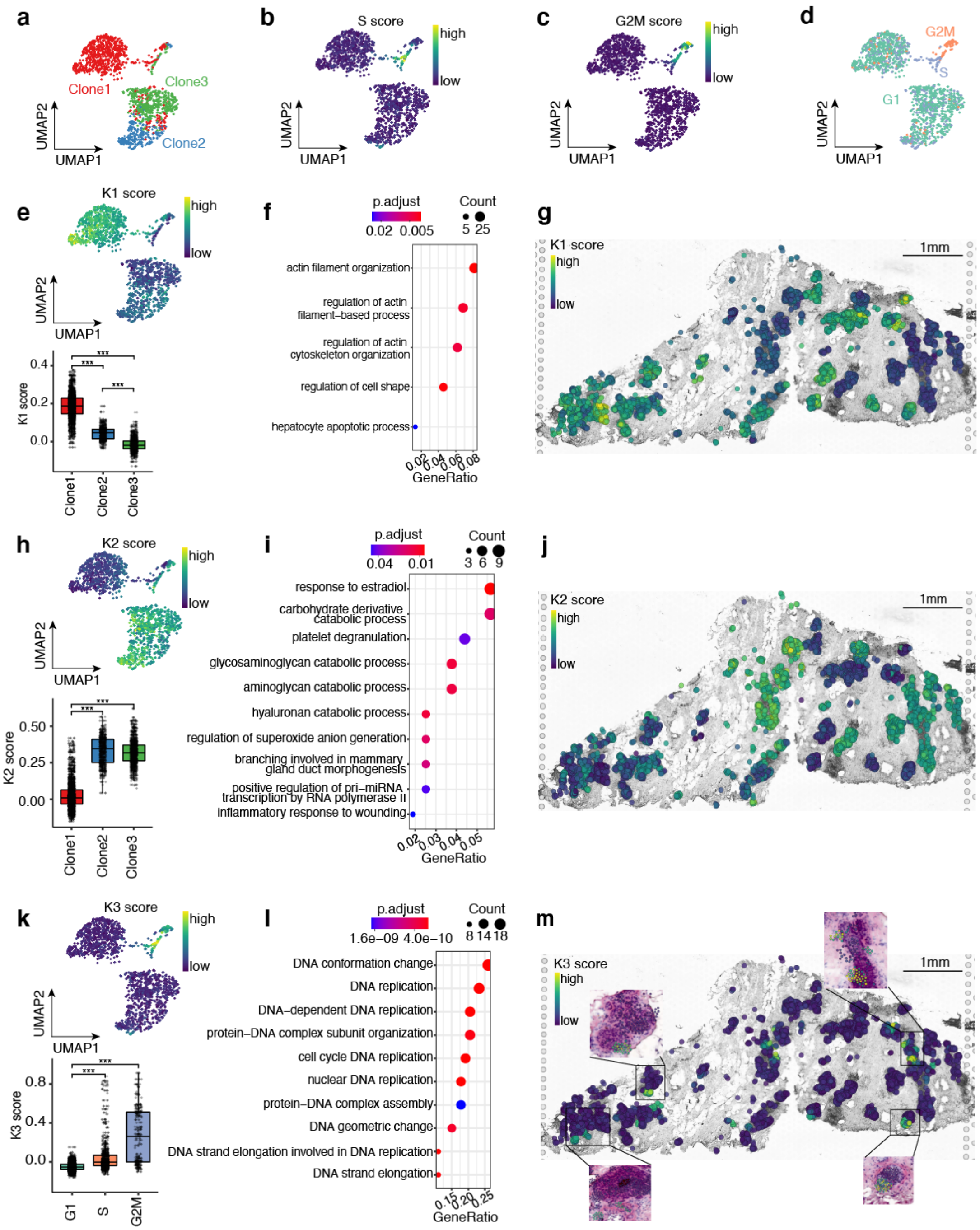


**Supplementary Figure 8 – Spatial co-expression patterns of tumor cells in DCIS1**

(**a**) Transcriptome-based UMAP of three tumor subclones identified from DCIS1 scRNA-seq data. Seurat-based cell cycle analysis on tumor cells showed S scores (**b**), G2M scores (**c**) and categorical cell cycle phases (**d**). (**e**) K1 module scores on the UMAP (upper) and boxplot across tumor subclones (lower). (**f**) GO enrichment analysis for the K1 module. (**g**) CellTrek map of K1 module scores. (**h**) K2 module scores on the UMAP (upper) and boxplot across tumor subclones (lower). (**i**) GO enrichment analysis for the K2 module. (**j**) CellTrek map of K2 module scores. (**k**) K3 module scores on the UMAP (upper) and boxplot across different cell cycle phases (lower). (**l**) GO enrichment analysis for the K3 module. (**m**) CellTrek map of K3 module scores. *** indicates $P < 0.001$ using Wilcoxon rank-sum test.

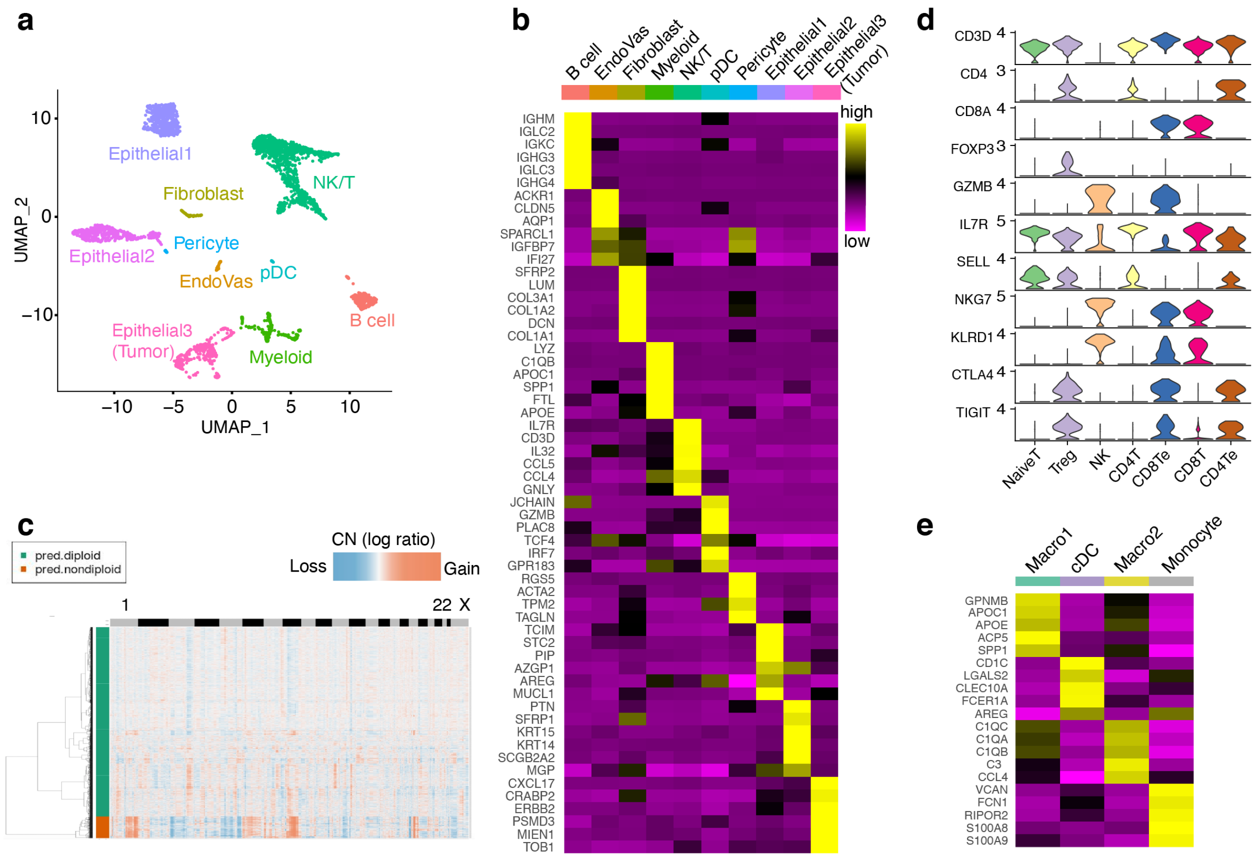


**Supplementary Figure 9 – ScRNA-seq data analysis of DCIS2**

(**a**) UMAP of DCIS2 scRNA-seq data, with clusters corresponding to different cell types. (**b**) Consensus heatmap of top differentially expressed genes for each cell cluster. (**c**) Heatmap of CopyKAT inferred copy number profiles from the scRNA-seq data and hierarchical clustering of cells. (**d**) Violin plots showing expression of canonical T cell state markers across different T cell states. (**e**) Consensus heatmap showing expression of top differentially expressed genes across different myeloid cell states.

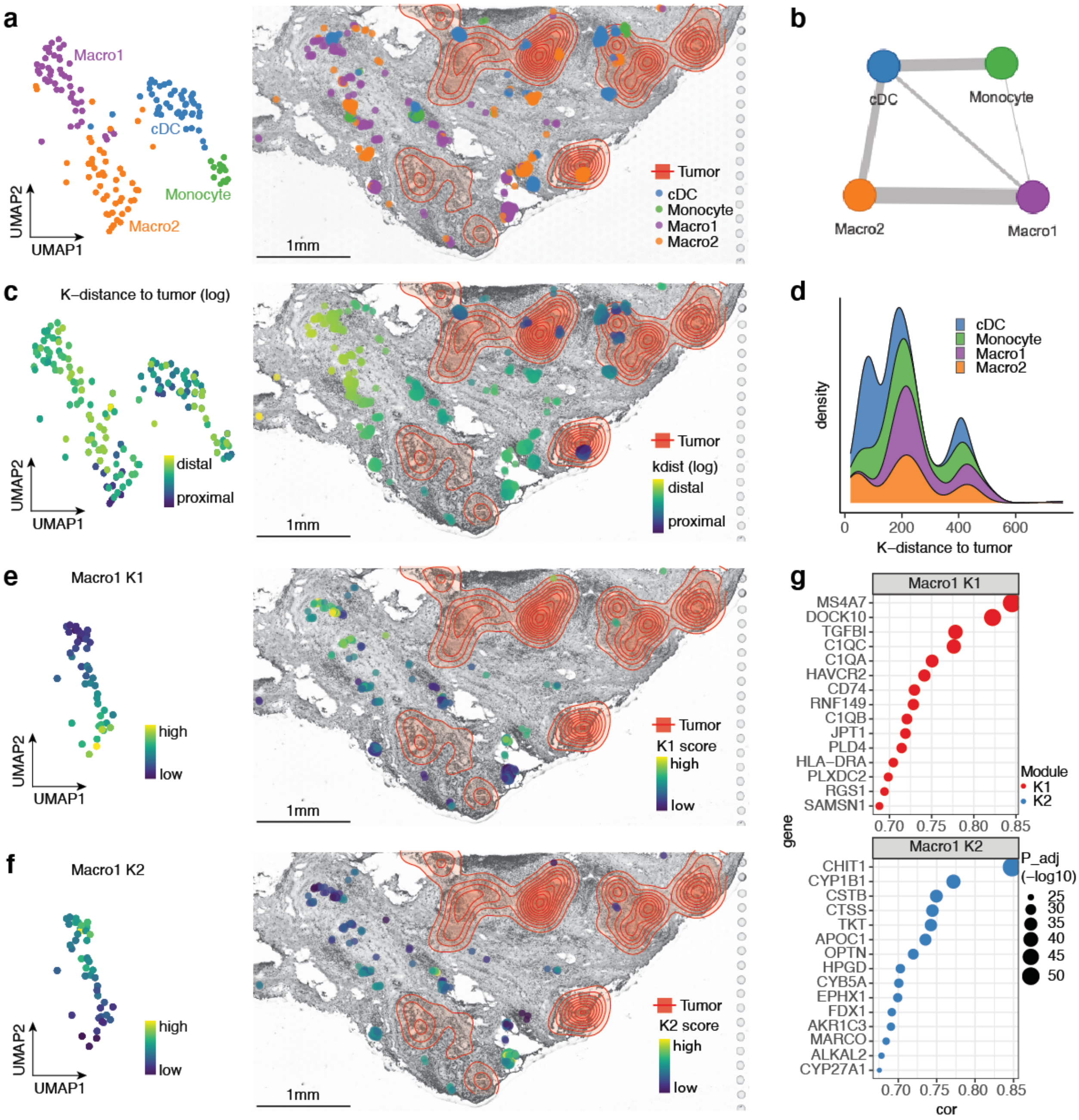


**Supplementary Figure 10 – CellTrek analyses of myeloid cell states in DCIS2**

(**a**) UMAP representation of myeloid cell states (left) and corresponding CellTrek map (right) (**b**) Spatial graph of myeloid cell states colocalization using SColoc. (**c**) UMAP (left) and CellTrek map (right) of myeloid cells K-distance to tumor cells. (**d**) Density plot of K-distances to tumor cells for each myeloid cell state. (**e**) UMAP (left) and CellTrek map (right) of Macro1 K1 module scores. (**f**) UMAP (left) and CellTrek map (right) of Macro1 K2 module scores. (**g**) Macro1 K1 (top) and K2 (bottom) module top correlated genes. The contour plots represent the densities of tumor cells.

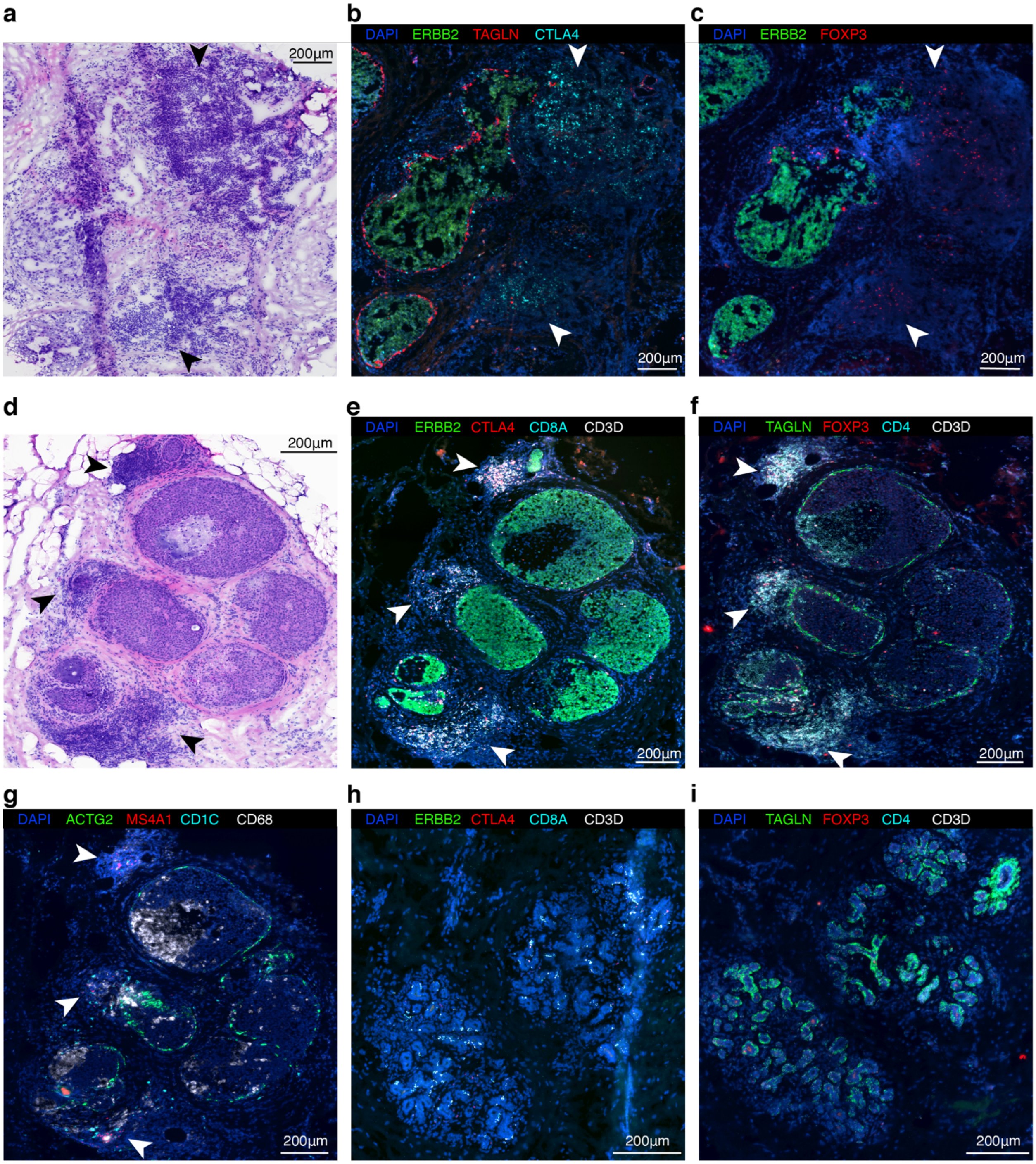
a
b
DAPI ERBB2 TAGLN CTLA4
c
DAPI ERBB2 FOXP3
d
e
DAPI ERBB2 CTLA4 CD8A CD3D
f
DAPI TAGLN FOXP3 CD4 CD3D
g
DAPI ACTG2 MS4A1 CD1C CD68
h
DAPI ERBB2 CTLA4 CD8A CD3D
i
DAPI TAGLN FOXP3 CD4 CD3D
200µm

**Supplementary Figure 11 – Orthogonal validation of tumor and immune cell spatial distributions using RNAscope assay in DCIS2 and DCIS3**

(a) H&E image of the DCIS2 tumor region. (b) RNAscope assay for *ERBB2, TAGLN* and *CTLA4* in DCIS2 tumor region. (c) RNAscope assay for *ERBB2* and *FOXP3* in DCIS2 tumor region. Data in (a-c) were collected from adjacent tissue slides. (d) H&E image of the DCIS3 tumor region. (e) RNAscope assay for *ERBB2*, *TAGLN* and *CTLA4* in DCIS3 tumor region. (f) RNAscope assay for *TAGLN*, *FOXP3*, *CD4* and *CD3D* in DCIS3 tumor region. (g) RNAscope assay for *ACTG2*, *MS4A1*, *CD1C* and *CD68* in DCIS3 tumor region. (h) RNAscope assay for *ERBB2*, *CTLA4*, *CD8A* and *CD3D* in DCIS3 adjacent normal lobular region. (i) RNAscope assay for *TAGLN*, *FOXP3*, *CD4* and *CD3D* in DCIS3 adjacent normal lobular region. Arrows point to different immune cell markers near the DCIS tumor ducts.

### Supplementary Tables

#### Supplementary Table 1 Parameter settings for different conditions in simulation-1

| | scRNA-seq | | | | ST | | | |
|---|---|---|---|---|---|---|---|---|
| Condition | Counts | Cell Proportion | Median Count | Median Gene | Counts | Spot Proportion | Median Count | Median Gene |
| 1 | 10 | 0.2 | 2815 | 494 | 50 | 0.2 | 15732 | 1526 |
| 2 | 10 | 0.4 | 2328 | 414 | 50 | 0.4 | 12516 | 1356 |
| 3 | 15 | 0.4 | 2195 | 414 | 75 | 0.4 | 12489 | 1356 |
| 4 | 15 | 0.5 | 1781 | 207 | 75 | 0.5 | 7006 | 338 |
| 5 | 20 | 0.5 | 1622 | 192.5 | 100 | 0.5 | 5055 | 320 |
| 6 | 20 | 0.6 | 929 | 63 | 100 | 0.6 | 2257 | 39 |
| 7 | 25 | 0.6 | 671 | 42 | 125 | 0.6 | 1537 | 23 |
| 8 | 25 | 0.7 | 444.5 | 31 | 125 | 0.7 | 1183 | 19 |

Counts means the UMI counts for each gene we downsampled and Cell/Spot Proportion means the proportion of cells/ST spots were used for downsampling. Median Count and Gene mean the median of UMI count and gene number after downsampling.

**Supplementary Table 2 Parameter settings for different conditions in simulation-2**

| Condition | Spatial Deviation |
|---|---|
| 1 | 30 |
| 2 | 60 |
| 3 | 90 |
| 4 | 120 |
| 5 | 150 |
| 6 | 180 |
| 7 | 210 |
| 8 | 240 |

Spatial Deviation means the deviation of the spatial gaussian noise we added to the cell.

**Supplementary Table 3 Parameter settings for different marker points in simulation-3**

| MarkerPoint | X-coordinate | Y-coordinate |
|---|---|---|
| 1 | 350 | 300 |
| 2 | 1300 | 1100 |
| 3 | 1950 | 800 |
| 4 | 1500 | 600 |
| 5 | 600 | 800 |
| 6 | 1000 | 1000 |
| 7 | 900 | 400 |
| 8 | 1700 | 1100 |

### Supplementary Data

#### Supplementary Data 1 – Screenshot of the CellTrek result for mouse brain data

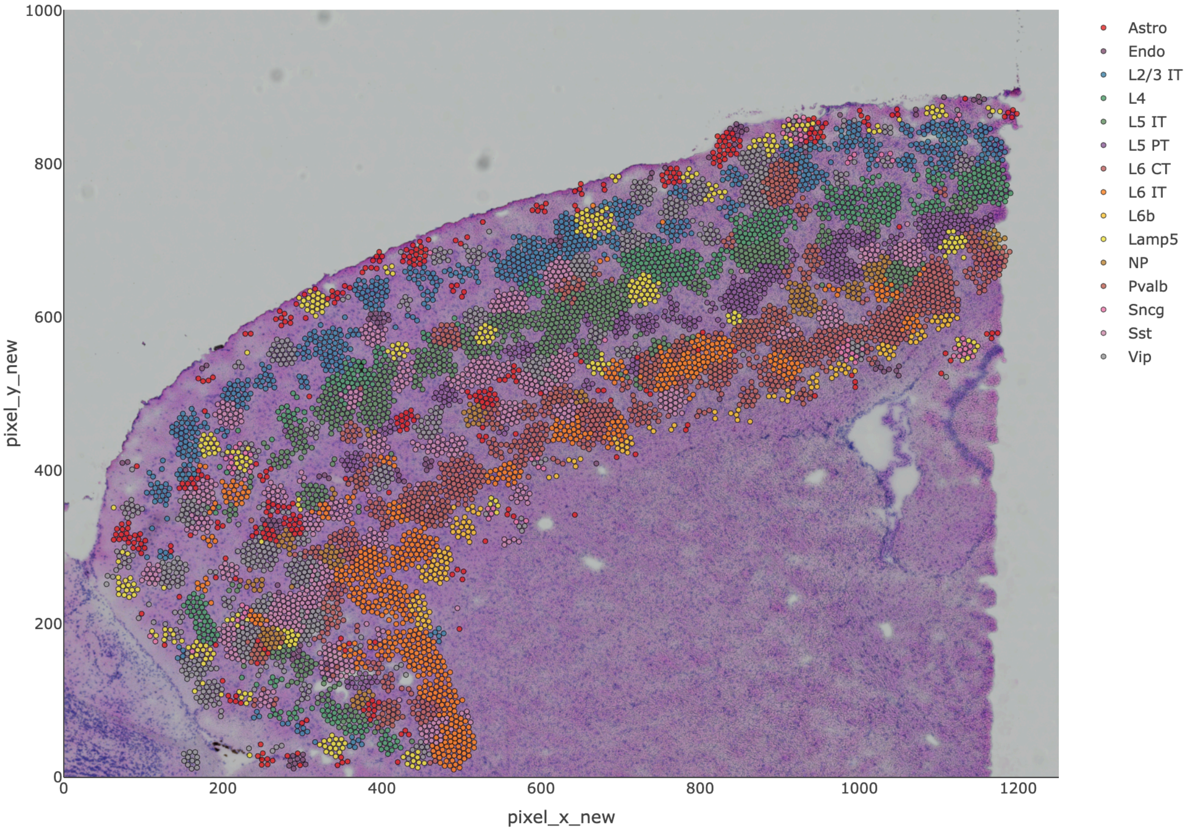

**Supplementary Data 2 – Screenshot of the CellTrek result for mouse kidney data**

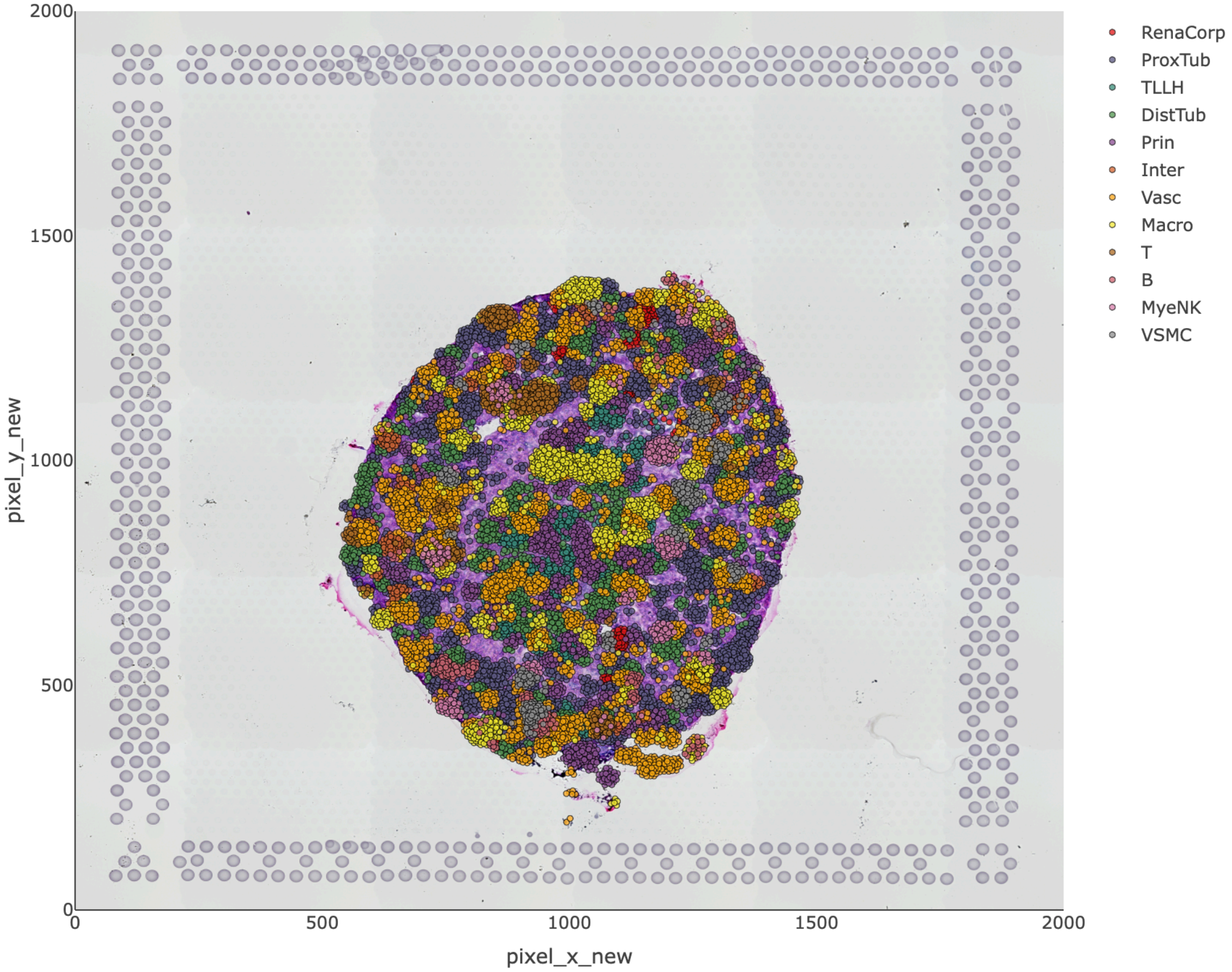

**Supplementary Data 3 – Screenshot of the CellTrek result for DCIS1 data**

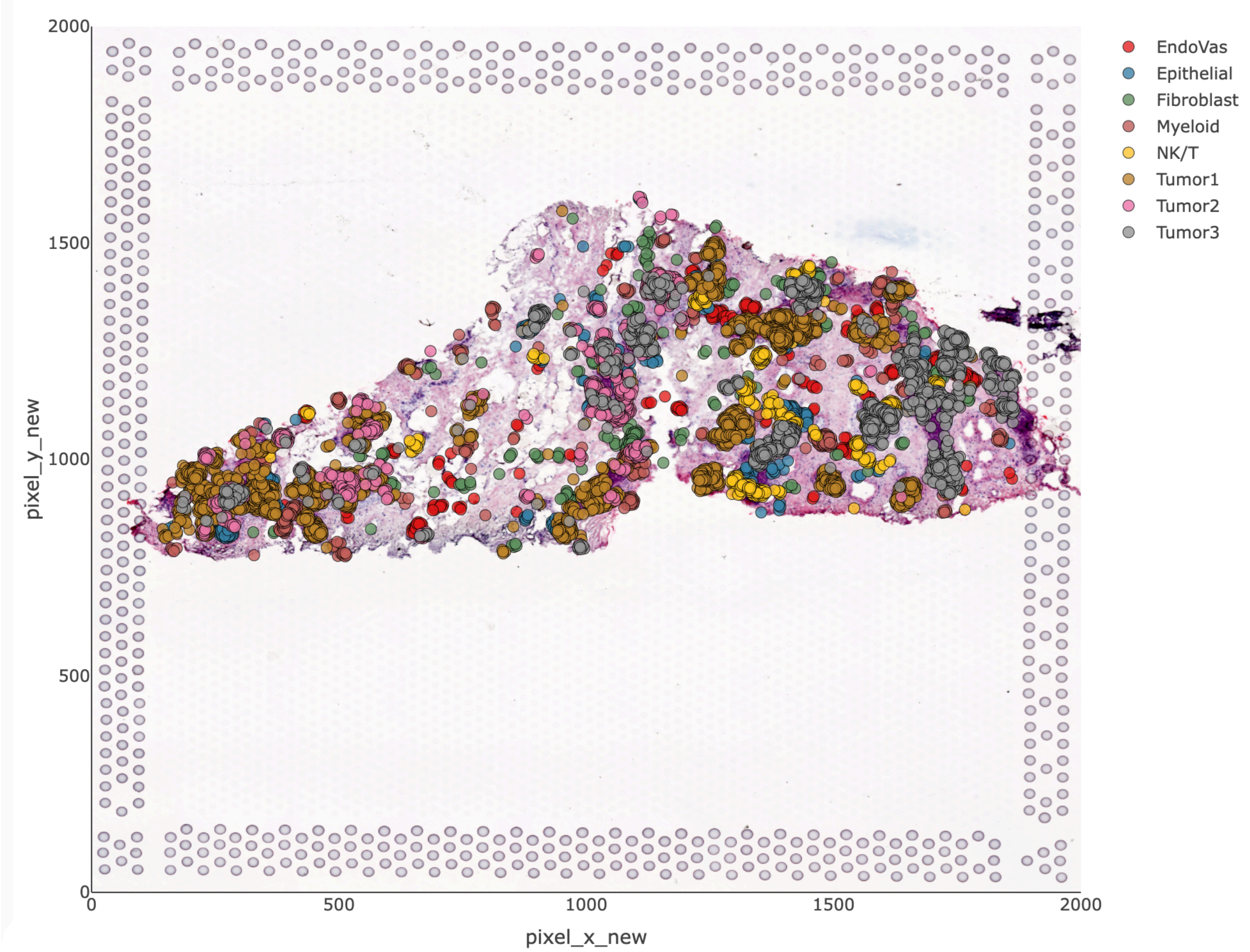

#### Supplementary Data 4 – Screenshot of the CellTrek result for DCIS2 data

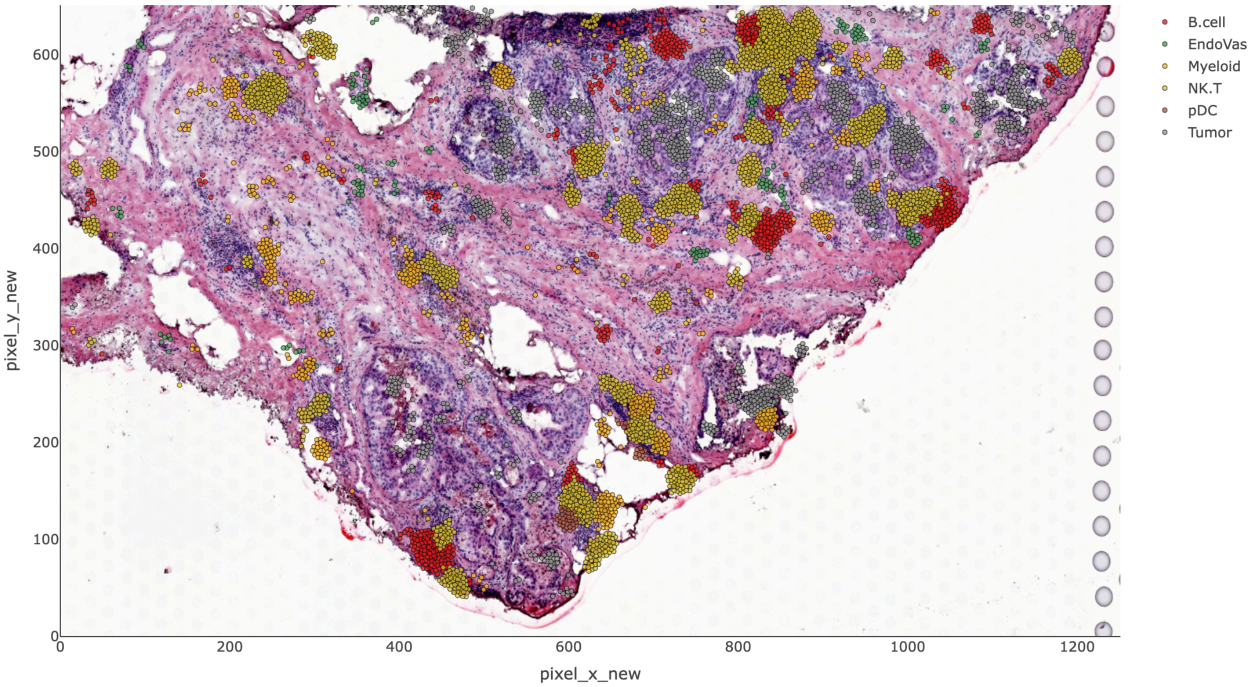
